# Correlative imaging of spatio-angular dynamics of molecular assemblies and cells with multimodal instant polarization microscope

**DOI:** 10.1101/2022.02.07.479448

**Authors:** Ivan E. Ivanov, Li-Hao Yeh, Juan A. Perez-Bermejo, Janie R. Byrum, James Y.S. Kim, Manuel D. Leonetti, Shalin B. Mehta

## Abstract

Biological function depends on the spatio-angular architecture of macromolecules - for example, functions of lipid membrane and cytoskeletal polymers arise from both the spatial and the angular organization of the constituent molecules. Correlative imaging of cellular and molecular architecture is valuable across cell biology and pathology. However, current live imaging methods primarily focus on spatial component of the architecture. Imaging the dynamic angular architecture of cells and organelles requires fast polarization-, depth-, and wavelength-diverse measurement of intrinsic optical properties and fluorophore concentration, but remains challenging with current designs. We report a multimodal instant polarization microscope (miPolScope) that combines a broadband polarization-resolved detector, automation, and reconstruction algorithms to enable label-free imaging of phase, retardance, and orientation, multiplexed with fluorescence imaging of concentration, anisotropy, and orientation of molecules at diffraction-limited resolution and high speed. miPolScope enabled multimodal imaging of myofibril architecture and contractile activity of beating cardiomyocytes, cell and organelle architecture of live HEK293T and U2OS cells, and density and anisotropy of white and grey matter of mouse brain tissue across the visible spectrum. We anticipate these developments in joint quantitative imaging of density and anisotropy to enable new studies in tissue pathology, mechanobiology, and imaging-based screens.

## Introduction

The spatial and angular distribution of proteins, lipids, nucleic acids, and carbohydrates is a key determinant, as well as informative readout, of biological function. Correlative live imaging of the architecture of cells and molecular assemblies is needed to visualize complex interactions among molecules and how these interactions determine functional states of cells. Imaging the spatial and angular distribution of biomolecules in terms of their density and anisotropy, respectively, can provide insights in how complex cell functions emerge. For example, emergence of contractile activity in the cardiomyocytes differentiated from the pluripotent cells can be analyzed over long term from their dynamic density and anisotropy, cellular impacts of perturbations can be profiled in terms of the density and anisotropy of organelles, and tissue components can be identified from spectral variations in their density and anisotropy. Making such correlative measurements require microscopes that enable multimodal imaging of spatio-angular architecture at high speed.

The spatial distribution of ensemble of biomolecules can be measured from changes in the optical path length they introduce using label-free quantitative phase imaging methods [1–3]. The spatial distribution of specific biomolecules can be measured via quantitative fluorescence imaging methods after labeling [4, 5]. In both label-free [6–8] and fluorescence modes [9–11], quantitative polarization microscopy enables measurement of anisotropy which reports the degree of alignment of assemblies of biomolecules and the angular organization below the spatial resolution limit of the microscope [9, 10, 12, 13]. Joint imaging of the spatio-angular distribution of unlabeled [14–17] and fluorescent [10, 13, 18–20] specimens with recently developed methods has enabled investigation of the architectural basis of cell and tissue function. Although substantial progress has been made in the design of optical systems and reconstruction algorithms for joint measurement of density and anisotropy, multiplexed imaging of many structures at timescales relevant to biology still remains challenging.

Measurement of density and anisotropy of multiple structures requires polarization, depth, and wavelength-diverse image acquisition, which remains slow with current designs. We have recently reported a method for combined quantitative label-free imaging of phase and polarization (QLIPP) and employed it for imaging the cell dynamics during mitosis and the architecture of brain and kidney tissue slices [16]. The method relies on illuminating the sample with light of different polarization states and acquiring through-focus images to reconstruct density, anisotropy, and orientation of cells and organelles. We have recently refined this method to illuminate the sample at different angles of incidence [17] and developed a reconstrution algorithm based on vector diffraction model, which allows us to measure the 3D spatio-angular architecture of the sample in terms of the distribution of its permittivity tensor. Using uniaxial permittivity tensor imaging (uPTI) we measured phase, principal retardance, 3D orientation, and optic sign of multiple organelles and structures in cells and tissues at high resolution. QLIPP and uPTI are readily compatible with other imaging modalities such as fluorescence or H&E staining and can be multiplexed into existing imaging pipelines [16, 17]. They are, however, limited in the acquisition speed they can achieve, as they sequentially illuminate the sample with light of different polarization or at different angles of incidence. In addition, these methods currently operate at a set wavelength due to the strong wavelength-dependent performance of many polarization optics. We have earlier reported an instantaneous fluorescence polarization microscope (instantaneous FluoPolScope) which can measure the position, intensity, anisotropy, and orientation of fluorophores from a single image. This is enabled by a detector with four polarization-resolved channels arrayed on the camera sensor such that the light is split by polarization state, rather than filtered by polarization state [10]. Using this technique, we studied the dynamics of actin and septin bundles in live cells. A similar image-splitting detection strategy was recently used in combining fluorescence polarization with STORM for super-resolution imaging of actin organization [13]. These methods provide insights in the spatio-angular architecture of organelles, but have a limited field of view as the four polarized channels are arrayed on a single detector. Taken together, imaging systems do not provide optimal imaging modes, resolution, and speed for correlative imaging of spatio-angular architecture in live cells.

Combining fluorescence and label-free polarization microscopy such that labeled molecular components can be studied in the context of the surrounding cell architecture is hampered by fundamental and experimental trade-offs. Label-free anisotropy measurements commonly rely on converting variations in polarization into intensities via filtering, i.e. using diattenuating optics which absorb non-transmitted states. Such detectors can have low detection efficiency for fluorophores, which makes them largely incompatible with fluorescence imaging. The performance of polarization optics, such as liquid crystal devices or polarizing beamsplitters, strongly depends on wavelength. As a result, anisotropy measurements have been constrained to a specific wavelength, preventing wavelength-based multiplexing of fluorescence and label-free imaging or imaging of multiple fluorescent targets on these instruments. In addition, combining fluorescence polarization and label-free imaging frequently requires changing the state of the microscope between the two imaging modalities. These changes are directed via software commands, which can introduce large delays in the acquisition pipeline.

Here we report multimodal instant polarization microscope (miPolScope) which enables multiplexing of fluorescence density, anisotropy, and orientation with its label-free counterparts – optical path length, retardance, and slow-axis orientation, at biologically-relevant timescales. We have balanced above design trade-offs as follows:

1. We have developed a broadband polarization-splitting detector that optimally uses available photons and provides high contrast polarization response across the visible spectrum.
2. We have implemented electronic sequencing of hardware states to minimize acquisition delays.
3. We have developed calibration and image analysis algorithms for quantitative imaging across large fields of view at improved signal-to-noise ratio.

We employed miPolScope to image architectural dynamics of HEK293T and U2OS cells simultaneously with fluorescently labeled actin network, plasma membrane, and DNA. We report 3D phase, retardance and slow-axis orientation measurements in beating cardiomyocytes at 10Hz. We also measure wavelength-dependent optical properties of grey and white matter in a mouse brain slice. These data demonstrate a performant computational imaging module that enables measurements of density and anisotropy across wavelengths, both in label-free and fluorescence modes, in live cells (see Visualizations 1 and 2).

## Results

### miPolScope’s light path and acquisition

miPolScope leverages a broadband instant polarization detector and microscope automation to multiplex label-free and fluorescence anisotropy imaging at multiple wavelengths. As with previous methods [8, 21] and as illustrated in fig. 1a, information about the sample anisotropy is encoded as differential intensities across polarization-resolved imaging channels (e.g. *1*0, *1*45, *1*90, *1*135). The density of structures (i.e., induced phase delay) and fluorophore concentration are measured from the propagation-dependent *XY Z* intensity distribution [9, 10, 16, 22]. miPolScope is designed as a module to enable easy integration with existing imaging systems (fig. S1), and can be attached to the camera port of commercial microscopes. This imaging module consists of a linear polarimeter which simultaneously detects light polarized along 0°, 45°, 90°, and 135° angles, similar to previous designs (fig. 1b) [8, 10, 13, 23, 24]. When constructing the four polarization-resolved imaging paths we selected optics (a non-polarizing 50/50 beamsplitter, two wire-grid polarizing beamsplitters, and an achromatic half-wave plate) with low wavefront distortions and relatively uniform polarization performance in the wavelength range of 430 - 770 nm (see methods and figs. S2 and 4 and table S1) to enable reliable calibration and analysis of the anisotropy of biological samples. For label-free imaging, the sample is illuminated in Köhler illumination with circularly polarized light, using a circular polarizer placed near the condenser aperture plane. For fluorescence imaging, the sample is illuminated with unpolarized light via a standard epifluorescence illuminator.

**Fig. 1.**
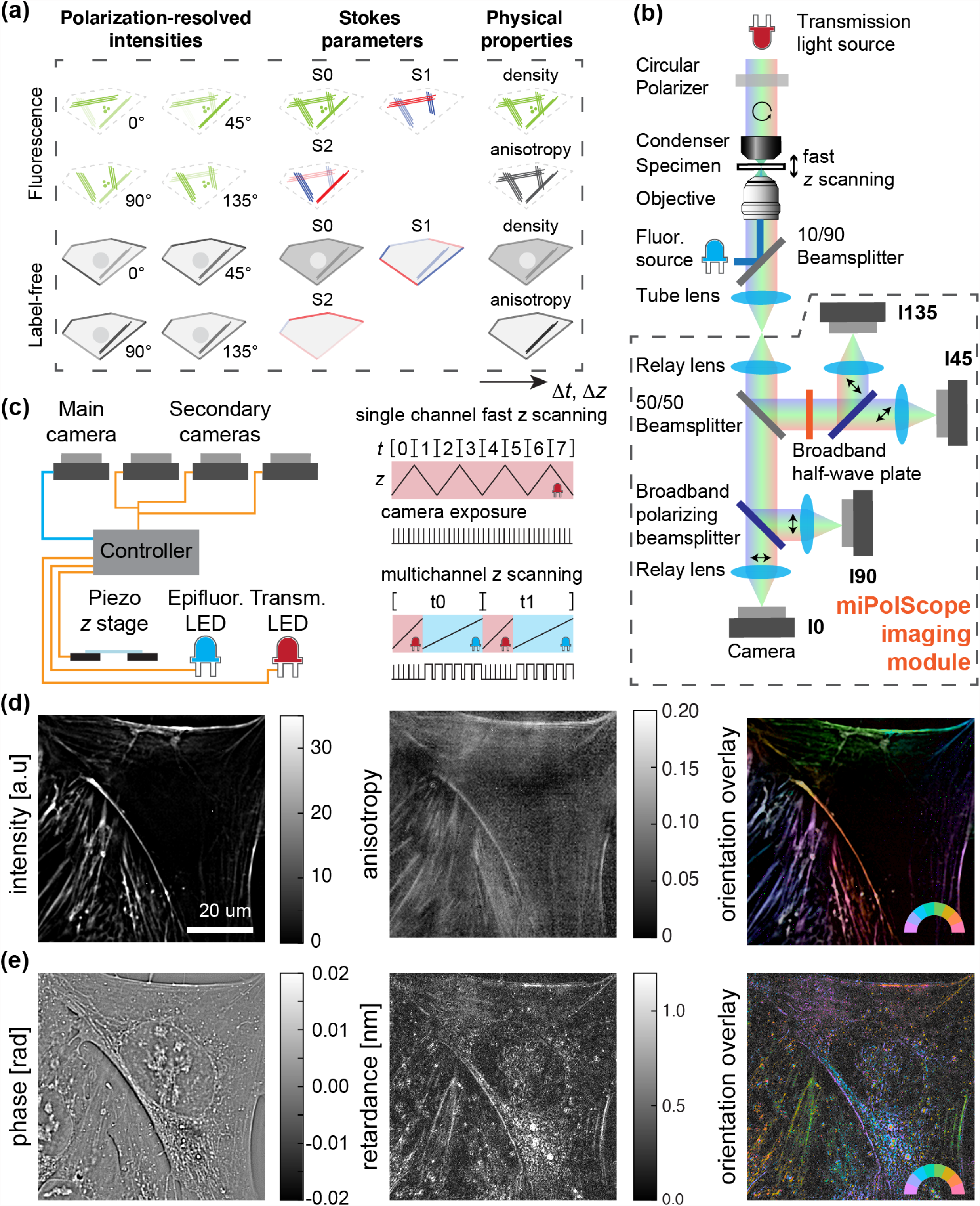
Fast measurement of density, anisotropy, and orientation in label-free and fluores-cence modes with miPolScope. **(a)** Illustration of the source of contrast in density and anisotropy measurements in label-free and fluorescence in 3D over time. **(b)** Schematic of the microscope. The miPolScope imaging module detects light linearly polarized along 0°, 45°, 90°, and 135°.**(c)** Fast imaging is enabled by automation of the microscope cameras, light sources, and piezo *z*-stage. **(d)** Deconvolved fluorescence intensity, anisotropy, and orientation measurements of live U2OS cells labeled with SiR-actin. An orientation colormap is given in the lower right corner. Brightness varies with fluorescence intensity and color saturation varies with anisotropy - isotropic structures appear white. **(e)** Phase, retardance, and orientation images of the same field of view as in **(d)**. Brightness varies with retardance and color saturation is fixed. See Visualizations 3 to 6 for full field-of-view time series and *z* fly-through movies.

miPolScope further leverages microscope automation (fig. 1c) to maximize the data acquisition rate, allowing us to capture fast dynamics in biological samples (discussed in fig. 3). The microscope is controlled through Micro-Manager [25, 26]. We use TTL pulse sequences to coordinate the acquisition by the cameras, movement of the piezo stage along *z*, and switching of the transmission and epifluorescence light sources using an Arduino Uno controller. In this “hardware sequence” acquisition mode, the cameras run continuously at a preset frame rate and the state of microscope components is changed via TTL pulses, eliminating delays due to software communication. Lastly, we drove the piezo stage in a custom triangle waveform when acquiring *z*-stacks at a rapid rate for a single channel (see methods). For multichannel imaging we increased the rate of data acquisition by using TTL triggering of the illumination sources in combination with quad-band filters, which eliminated the need for switching filter cubes.

We employ the Stokes formalism [10, 16, 17] to calibrate the polarization response of the light path which is represented by an “instrument tensor” *A* (discussed in fig. 2). Measured raw intensities are converted into Stokes images by multiplying with the inverse instrument tensor and these Stokes images are then used to reconstruct the sample physical properties. We employ background correction and image denoising as described in fig. 2 to enhance the signal from weakly birefringent biological samples. We provide Jupyter notebooks demonstrating steps of the data reconstruction pipeline at https://github.com/mehta-lab/miPolScope.

**Fig. 2.**
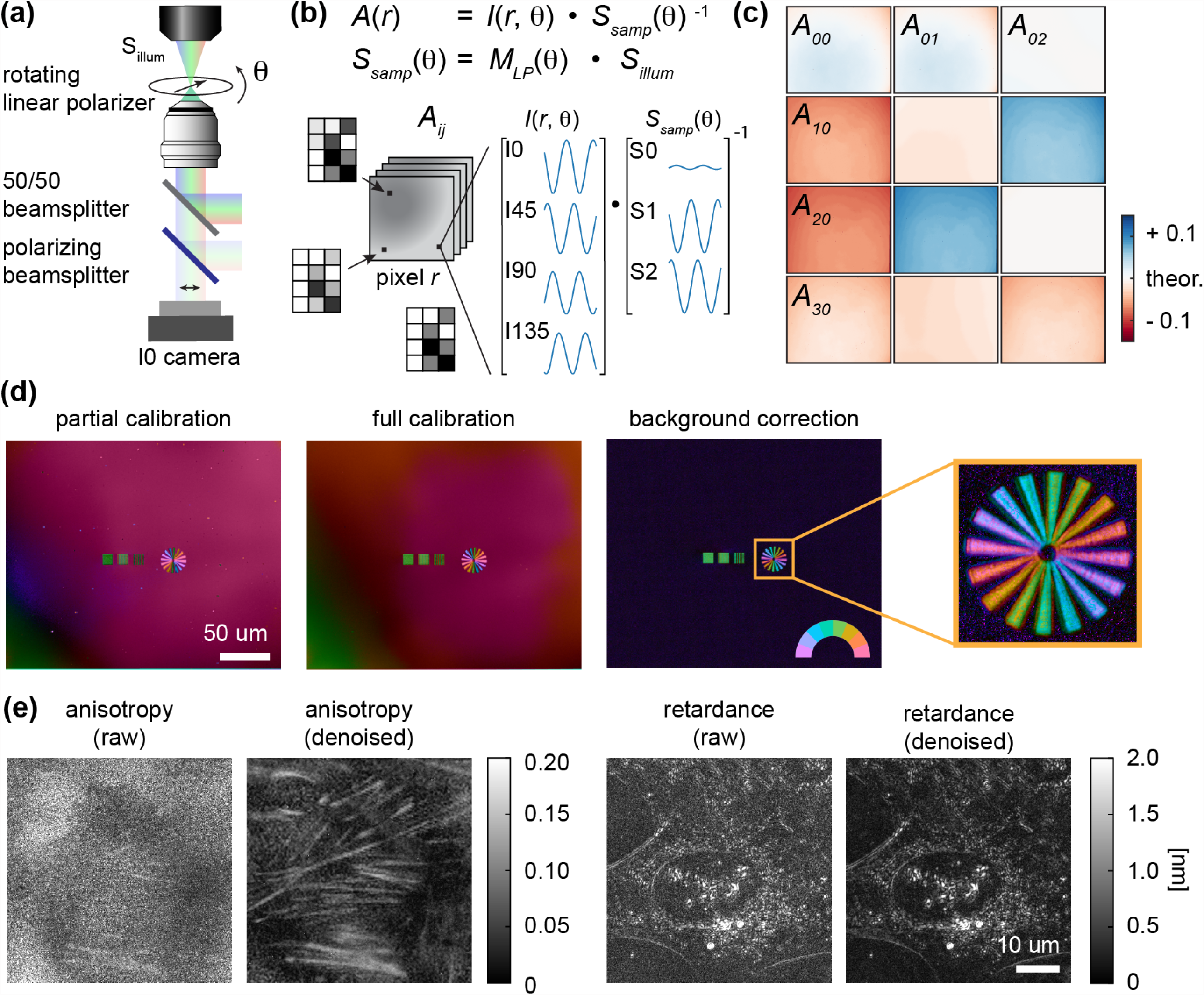
New calibration method and data denoising improve reconstruction of anisotropy. **(a)** The spatially-resolved instrument tensor *A*_*ij*_(*r)* of the miPolScope detector is calibrated by illuminating at the imaging wavelength and rotating a linear polarizer in the sample plane to generate a variety of polarization states (see methods). **(b)** *A (r)* is calculated from intensities at every pixel *r, 1(r)*, and Stokes states of light in the sample plane, *S*_*samp*_*(θ*)= *M*_*LP*_ *(θ*)*S*_*illum*_, measured at varying polarizer angles *θ*. **(c)** Images of components of the instrument tensor *A*_*i j*_. (d).Reconstruction of an anisotropic glass target using the full calibration procedure described here reduces polarization aberrations compared to a reconstruction using a partial calibration procedure, which omits calibration of *S*_*illum*_ and uses an average instrument matrix. Background correction further removes birefringence signal originating from the sample substrate. Zoom-in confirms that the slow-axis orientation of the anisotropic glass target is measured as expected from its design. (e).Denoising improves fluorescence anisotropy and label-free retardance measurements. See visualizations 7 and 8 for side-by-side comparison in time series movies.

With these technological advances, we were able to carry out measurements of fluorescence density, anisotropy, and orientation multiplexed with label-free measurement of cell density, anisotropy, and orientation which together report the architecture of organelles in the context of the architecture of the cell. Figure 1d-e and visualizations 3 to 6 show fluorescently labeled actin bundles in U2OS cells. We observe aligned stress fibers near the basal membrane of the cell. The measured SiR-actin ensemble dipole orientation is perpendicular to the fibers, in agreement with previous measurements [27, 28]. Phase measurements show the cell membrane, nucleoli, and vesicles; phase variations within the nucleolus are also observed, consistent with different compartments within this phase-separated organelle [29, 30]. Retardance and orientation measurements show boundaries of the cell and organelles, and thick cytoskeletal filaments. Relative to previous methods, miPolScope allows fast measurements of density and anisotropy in dynamic samples (fig. 3), integration of anisotropy measurements of multiple fluorophores with label-free imaging at different wavelengths (fig. 1d-e, fig. 4a-b), and further enables measurements of physical properties of biological samples across the spectrum (fig. 4c-d).

**Fig. 3.**
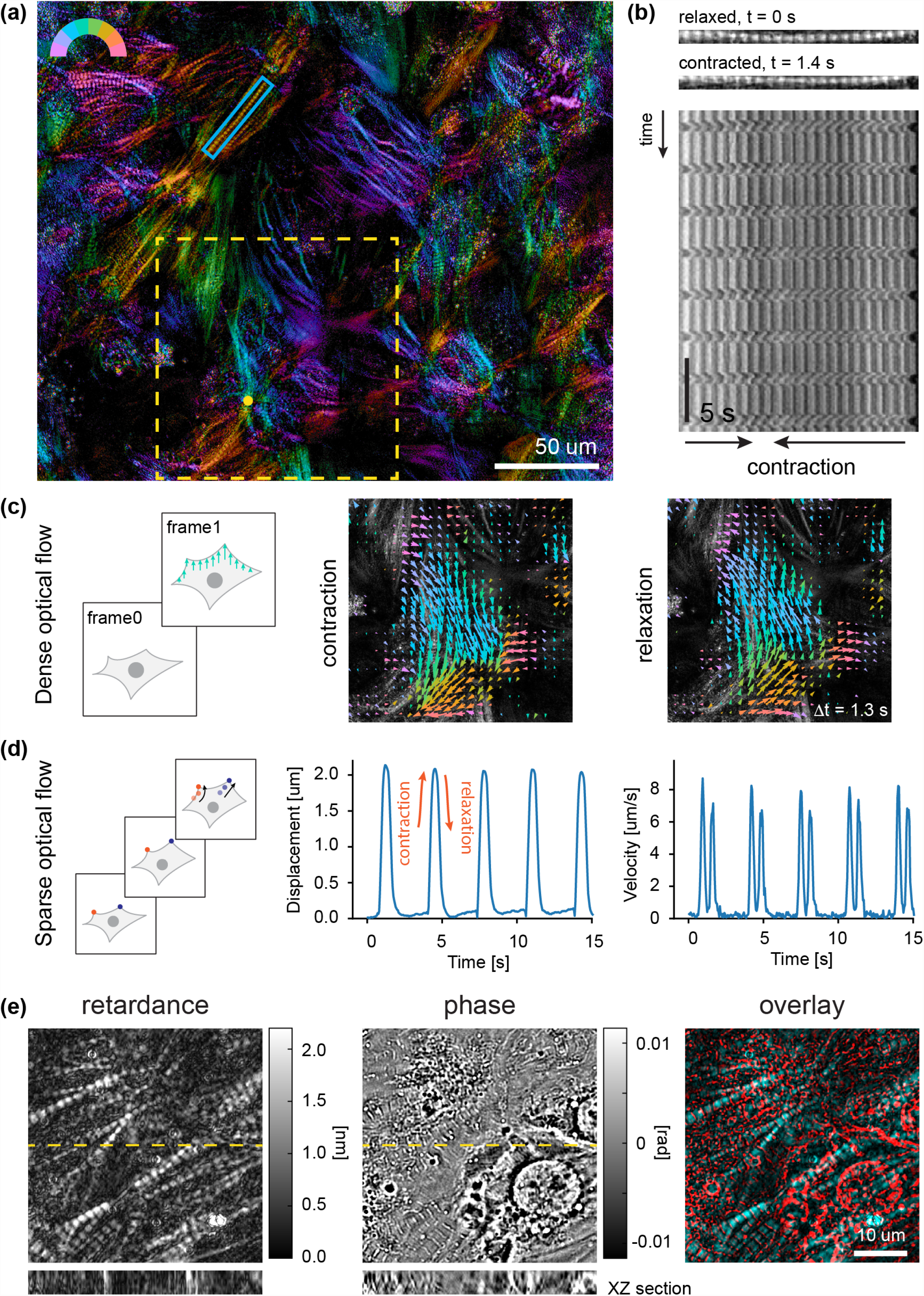
Label-free imaging of sarcomere dynamics in live cardiomyocytes. **(a)** Retardance and orientation overlay of iPSC-derived cardiomyocytes in 20% OptiPrep index matching media. Blue rectangle, yellow rectangle, and yellow dot show regions used for analysis in panels **(b), (d)**, and **(e)**, respectively. **(b)** Kymograph along a single myofibril. Retardance images at the top show sarcomere in relaxed and in contracted states. **(c)** Dense optical flow analysis of myofibril dynamics. Arrows are shown using the same colormap as in **(a)** and 5x longer than the measured displacement. **(d)** Displacement and velocity measurements from sparse optical flow analysis. **(e)** Combined retardance and phase imaging of beating cardiomyocytes; *XZ* cross-sections through the center of the image (dashed yellow lines) are shown underneath. See visualizations 9 to 13 for associated movies.

**Fig. 4.**
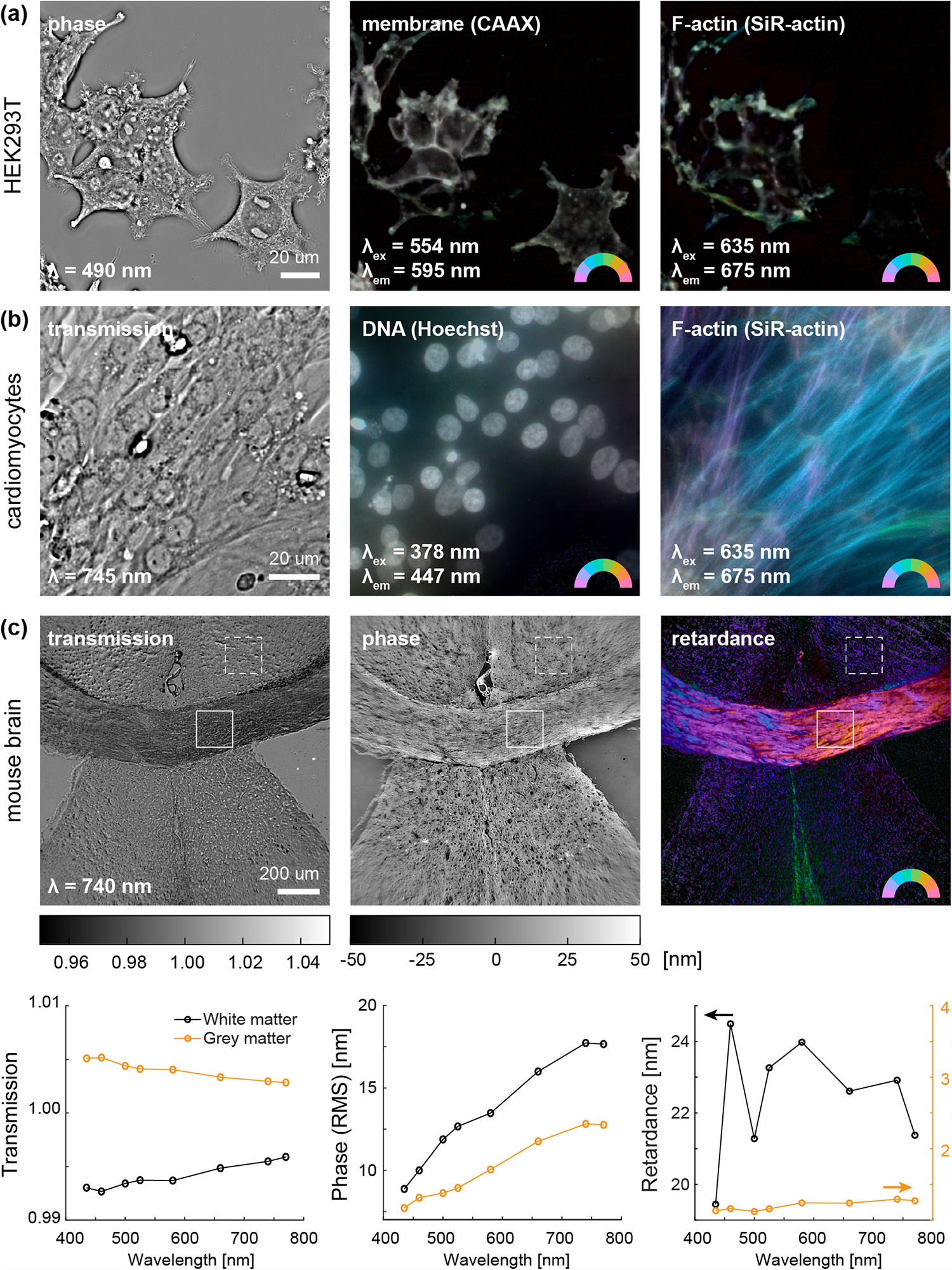
Multichannel imaging of density and anisotropy across the visible spectrum. **(a)** Multiplexed imaging of phase and fluorescence anisotropy of plasma membrane labeled with CAAX-mScarlet and F-actin stained with SiR-actin in HEK293T cells. **(b)** Multiplexed imaging of transmission and fluorescence anisotropy of DNA stained with Hoechst-33342 and F-actin in iPSC-derived cardiomyocytes. See visualizations 14 to 16 for associated movies. **(c)** Transmission, phase, and retardance imaging at 740 nm of a mouse brain slice around corpus callosum. White solid and dashed boxes show regions within the white and grey matter, respectively, used for spectral analysis below. Note second y-axis used for grey matter retardance in bottom right plot.

### Reconstruction of density and anisotropy

Quantitative data reconstruction requires calibration of the microscope and correction of background introduced by the sample chamber, as we have previously described [10, 16, 17]. Here we extend these methods to address challenges inherent in the design of the miPolScope detector.

The parallel polarization detectors of the miPolScope imaging module enable anisotropy measurements in the imaging plane at the acquisition rate of the cameras, >10x faster than other state-of-the-art quantitative phase and polarization imaging methods, such as QLIPP [16], which sequentially excite the sample with light of different polarization. This increase in acquisition speed comes with increased isotropic background, which leads to higher noise in the anisotropy measurements. QLIPP uses elliptically polarized excitation light and a circular analyzer, which suppresses the isotropic background and associated photon shot noise in the recorded images. miPolScope, on the other hand, uses circularly polarized excitation and linear detectors, which leads to decreased signal-to-background ratio in the raw image. The use of four independent cameras further increases the attainable field of view relative to previous methods [10, 13], up to the field number supported by the microscope. Imaging a large field of view leads to propagation of light at large angles in the aperture plane. Polarization optics commonly exhibit an angle-dependent response, and when placed in the aperture plane can introduce background in the image space. To enable quantitative measurements of biological samples, which commonly exhibit retardance on the order of λ,/100 or less, we developed a calibration procedure which accounts for position-dependent polarization response in sample space, implemented background correction strategies to compensate for birefringence introduced by the sample substrate, and employed wavelet-based denoising to improve the SNR in anisotropy measurements (fig. 2).

### Calibration

The detection path of miPolScope is calibrated by transmitting light of known Stokes states of polarization through the microscope and measuring the resulting intensity. An “instrument matrix” is then computed, which transforms the known input polarization state into intensity measurements [10, 17, 31, 32]. During data reconstruction, the inverse of the instrument matrix is used to determine how the sample perturbs the polarization state of the illumination light based on the recorded intensity images. Previously we averaged the intensity of the recorded images before computing the instrument matrix [16, 17]. Here we extend this approach by calibrating the microscope for every pixel in image space to compute an instrument tensor, which now also accounts for position-dependent polarization distortions over the field of view (fig. 2a-c). The calibration process consists of rotating a linear polarizer in the sample plane and recording intensity images on the four detectors of the miPolScope imaging module (fig. 2a), which are registered in space (fig. S3 and table S2). In this process, we calibrate the Stokes state of the illumination light source, *S*_*illum*_, and the instrument tensor, *A* (fig. 2b, fig. S4, and methods). A unique calibration is carried out at each label-free imaging wavelength or fluorophore emission wavelength. Calibration of the detection path for fluorescence anisotropy measurements is carried out in the same manner, except that the microscope is illuminated in transmission at the fluorophore emission wavelength. Figure 2c shows calibrated components of the instrument tensor. The calibration accounts for deviations from the theoretical values of the instrument matrix for an ideal linear polarimeter, as well as variations across the image dimensions due to position-dependent polarization distortions.

### Background correction

Data reconstruction using the instrument tensor corrects for large-scale background introduced by polarization optics, as well as small-scale image blemishes such as dust. Figure 2d shows a comparison of data reconstruction of a birefringent test target [33] using the full calibration procedure outlined above and a partial calibration which uses an instrument matrix (i.e. instrument tensor averaged over the image dimensions) and assumes that the illumination source is circularly polarized. Reconstruction based on the full calibration procedure reduces bias in the measured retardance and orientation. The sample substrate contributes significantly to the observed background as well. Background correction using an empty field of view removes reconstruction bias and reveals that the measured orientation in the spokes of the test target follows their morphological orientation, as designed.

### Denoising

The miPolScope imaging module enables light-efficient detection with polarizing beamsplitters (fig. 1b and fig. S1), but they also lead to higher transmission of the circularly polarized excitation light, and consequently higher photon shot noise. Biological structures often have low retardance which leads to small intensity modulation in the detection channels that can be obscured by noise. We leveraged image denoising through *z*-projection and wavelet filtering (see methods) to remove uncorrelated intensity fluctuations and enhance the SNR in anisotropy images. Figure 2e and visualizations 7 and 8 show a comparison of fluorescence anisotropy and retardance computed from raw and denoised data. Denoised images enable easy interpretation and quantitative measurements.

The reconstruction pipelines for fluorescence and label-free data, consisting of Stokes image computation, background correction, and denoising, are illustrated with examples in figs. S5 and S6.

### Fast label-free imaging of cellular activity

Instant measurements of anisotropy, combined with hardware sequencing, enable tracking of the dynamics of cells and organelles at high spatial and temporal resolution. Cardiomyocytes derived from induced pluripotent stem cell (iPSC) have emerged as a model system for studying cardiac diseases and for drug screening [34–36]. Their characteristic contractile dynamics are informative of cellular state and functional output, but are commonly difficult to study due to their high beat frequency of 20 - 100 beats per minute [37, 38]. Using miPolScope, we overcame limitations in the data acquisition rate, and collected high-speed measurements of myofibril architecture and activity in spontaneously beating cardiomyocytes (fig. 3).

Figure 3a shows a retardance and orientation overlay from a movie of beating cardiomyocytes acquired at 20 Hz at full field of view of the microscope. A network of myofibrils is visible with the anisotropic A-band, which overlaps with the length of the myosin thick filament, clearly resolved. The slow axis orientation, encoded in color, follows the direction of the filaments. Visualization 9 shows a side-by-side comparison of computed transmission and overlay of retardance and orientation over time. Cell nuclei, which are visible in the transmission image, indicate that the observed sarcomere network spans across multinucleated cells.

In our initial experiments, the visibility of sarcomeres was masked by edge contrast introduced by the crowded cytoplasm (fig. S7). Density variations in the cytoplasm introduce edge birefringence, which can mask form birefringence of aligned structures. For imaging, we placed the cells in media with refractive index tuned to reduce phase variations and edge birefringence (see fig. S7 and methods) [39]. This approach allowed us to visualize fine substructure of sarcomeres, including A-band, I-band, and z-disk (fig. 3e).

We zoom in on an individual myofibril in fig. 3b and show its movement over time as a kymograph of the retardance channel. We can track the position of sarcomeres as a function of time and we observe that the fiber contracts asymmetrically, with the region of least displacement shifted to the left. We also performed dense and sparse optical flow on a sub-region of the field of view (fig. 3c-d and visualization 10). Dense optical flow is useful for tracking the velocity magnitude and direction over a given region, whereas sparse optical flow is useful for tracking the position of specific features over time (schematics on the left) [40, 41]. Figure 3c shows the displacement within a region of interest over one cycle of contraction and relaxation. The color of the arrows encodes the direction of motion and generally aligns with the morphological orientation of the myofibrils and the measured slow axis orientation. Figure 3d shows the displacement and velocity trajectories of a point highlighted in panel **(a)**. We measured beat frequency of 18 beats per minute, in accordance with previous results [37, 38, 42, 43].

The custom triangle *z*-axis waveform we used for moving the piezo stage (fig. 1c) allows us to rapidly acquire 3D volumes in a single channel. We used this method to collect 3D label-free data at a rate of 1 volume (20 images) per 100 ms and reconstruct phase in addition to retardance and orientation. As the data acquisition rate of the miPolScope is limited by the sustained camera framerate, we had to reduce the field of view to acquire images at 200 Hz. Figure 3e shows snapshots of retardance, phase, and phase-and-retardance overlay for a sub-region of the field of view. Visualizations 11 and 12 show a time series movie and *z*-stack fly-through of the full dataset and visualization 13 shows dense optical flow analysis. For this sample we measured beat frequency of 110 beats per minute, consistent with an earlier stage post differentiation compared to cells in fig. 3a. Phase imaging reveals the protein-dense z-disk in sarcomeres, as we previously reported for fixed samples [17]. Label-free polarization imaging provides quantitative structural information not accessible with conventional brightfield or fluorescence microscopy, and can be a powerful tool for following the long-term progression of cardiac disorders in iPSC models or studying the pathology of unstained cardiac specimens.

### Multimodal imaging of organelle, cell, and tissue architecture

We demonstrate multimodal live cell imaging, reporting on the overall cell architecture in phase or transmission, multiplexed with molecular labels for DNA, F-actin, or membrane. Such quantitative measurements can reveal new connections between the architecture of cells and constituent organelles.

Figure 4a shows snapshots from a movie of HEK293T cells exploring their surroundings. The cells express CAAX-mScarlet, which localizes to the plasma membrane, and were labeled with SiR-actin to stain actin structures. We acquired z-stacks at a rate of one multichannel dataset per 45 seconds to observe changes in the cell morphology over time (see visualizations 14 and 15). The phase channel reports on quantitative density changes within the cell, and can be used to track the overall cell morphology and health, or the dynamics of specific organelles, such as the nucleus, nucleolus, vesicles, or cell protrusions. Fluorescence anisotropy measurements show that CAAX-mScarlett localizes to the plasma membrane; we detect isotropic angular distribution (indicated by the white color in the image), as the mScarlett fluorophore is connected to the CAAX peptide via a flexible linker and is free to rotate. SiR-actin labeling shows primarily anisotropic cortical actin structures.

Figure 4b shows beating cardiomyocytes in transmission, and Hoechst-33342 and SiR-actin staining, as another example of multimodal live imaging. The three channels were acquired sequentially to maximize the data acquisition rate for each channel (see visualization 16). The transmission image reports on the overall morphology of the cell layer, whereas the two fluorescent labels report on individual cell markers (the nucleus) and extended actin network spanning multiple cells. SiR-actin-stained filaments are anisotropic with orientation perpendicular to the filament, as in previous examples. The DNA stain appears isotropic as multiple fluorophores, each intercalated into DNA at different orientation, contribute to the signal at each voxel.

The broadband performance of the miPolScope imaging module further enables label-free measurements of structures at multiple wavelengths. Different materials are expected to show characteristic variations in physical properties derived from label-free measurements, which can be used to identify specific structures and disambiguate label-free data [44–46]. Figure 4c shows label-free measurements and analysis at 8 distinct wavelengths of grey and white matter near the corpus callosum in a mouse brain slice. Transmission measured relative to a background region shows that white matter becomes more optically opaque with increasing wavelength, whereas grey matter becomes more optically transparent. Optical path length, as reported by the root mean squared (RMS) phase value, increases with wavelength for both grey and white matter, at different rates. Retardance of the white matter shows variation with wavelength, while retardance in the grey matter remains low and relatively constant with wavelength. This analysis can be extended to multiple materials and structures, and combined with machine learning models [16, 47] to identify features of interest in biological samples.

## Discussion

Multimodal instant PolScope (miPolScope) exploits polarization, depth, and wavelength diversity to encode the density and anisotropy intrinsic to cell structures or reported by a fluorescent label into images. These images are calibrated, background corrected, denoised, and reconstructed to provide high resolution imaging of the spatio-angular architecture of organelles, cells and tissues. The miPolScope imaging module is easy to integrate into many microscopes. We have demonstrated multiple applications of the method, including live imaging of architectural dynamics in beating cardiomyocytes, tracking of labeled molecular components in HEK293T and U2OS cell lines, and detection of changes in optical density and anisotropy of grey and white matter in a mouse brain slice as a function of wavelength.

The capability of imaging spatio-angular architecture in multiple channels can provide new correlative measurements that provide insights in mechanobiology of cells, e.g., how the ordered arrangement of myofibrils establishes mechanical linkage across cardiac cells or how the labeled F-actin network regulates the shape of the plasma memebrane visualized without the label.

Physical properties and morphological features of cellular organelles can carry enough information to uniquely identify them in label-free images. In the case of sarcomeres in cardiomyocytes, miPolScope provided measurements of dynamic architecture that otherwise require substantial optimization of fluorescent labeling. Unlike fluorescence, label-free imaging is not phototoxic to cells or limited in duration by photobleaching, and can be used with specimens not amenable to live staining, such as primary tissue samples. Quantitative label-free measurements can further reveal subtle architectural alterations, before larger-scale disease-relevant morphological changes occur. miPolScope is well suited to long-term studies of the dynamics of cardiomyocytes in disease model systems or in high-throughput drug screens.

The design of the miPolScope imaging module has been optimized for combined anisotropy measurements in fluorescence and label-free contrasts. Fluorescent dipoles emit linearly polarized light, and thus a linear polarimeter based on polarizing beamsplitters is an optimal choice for fluorescence anisotropy measurements. However, label-free imaging with circularly polarized illumination and linearly polarized detection is sub-optimal due to the transmission of isotropic background light, whose shot noise (fig. 2) can mask intensity modulations due to specimen anisotropy. Sensitive label-free imaging has been achieved using illumination with elliptically polarized light and detection through a circular analyzer of opposite handedness [8, 16]. A future design of multi-path polarimeter with tunable detection states based on fast programmable liquid crystal devices may provide anisotropy measurements with optimal SNR in fluorescence and label-free contrasts.

Combined spectral and polarization imaging is an emerging area of research with sensors and techniques being developed to provide more complete measurements of the properties of light [44, 48–52]. These methods are widely used for analyzing tissue pathology, for example, identifying skin lesions [53] or detecting residual cancerous tumor after treatment [54]. Using miPolScope, we demonstrated density and anisotropy measurements at eight different wavelengths in distinct regions of the mouse brain. Spectral label-free measurements with miPolScope may enable analysis of the tissue pathology as well as discovery of cell types and their distribution in unstained specimens.

Current design of miPolScope projects angular anisotropy of the specimen onto the focal plane and is insensitive to anisotropy out of the focal plane, similar to QLIPP [16] and related designs. This is not a significant limitation when imaging cells that are typically flat, but can be a limiting factor when imaging 3D spatio-angular architecture of tissues. 3D anisotropy of live specimens can be measured by refining the angle-resolved imaging used in uPTI [17] and combining it with the fast acquisition reported here.

The measurements reported here rely on single-photon interactions of the specimen with light - fluorescence emission or light scattering, which are well suited to live cell imaging due to high resolution and compatibility with widely used fluorescent markers. Single photon imaging is, however, limited in penetration depth and commonly useful in tissues up to 150 µm thick, especially when combined with silicone immersion objectives that minimize spherical aberrations. Nonlinear multiphoton light-matter interactions such as two-photon fluorescence or second harmonic generation enable label-free imaging of autofluorescent molecules (e.g. nicotinamide adenine dinucleotide (NADH), flavin adenine dinucleotide (FAD)) or anisotropic structures (e.g. collagen, myosin thick filaments, microtubules) [55]. Such methods use near-IR excitation, penetrate deeper into tissues, and can be multiplexed to enable imaging of cells and tissues with prominent applications in cancer detection [56–58]. The concepts and algorithms reported for miPolScope can be extended to multiphoton imaging techniques to provide a more complete view of tissue morphology and pathology.

In conclusion, miPolScope achieves polarization-, depth-, and wavelength-diverse imaging, and enables measurement of density and anisotropy of biological structures with or without label. Multiplexed measurements can be made across the visible spectrum and at high temporal and spatial resolution. We anticipate that the high spatio-temporal resolution and quantitative modes of information of miPolScope will prove valuable for drug screens, as well as studies in mechanobiology, tissue pathology, and systems biology.

## Materials and Methods

### Microscope layout

The microscope is constructed using a RAMM frame from Applied Scientific Instrumentation and uses a Nikon Plan Apo 60x 1.20 NA water immersion objective. The miPolScope imaging module (fig. 1b) uses a TTL200-A lens and four TTL100-A lenses (Thorlabs, Inc.) to re-image the primary image plane at 0.5x magnification onto four separate acA2440-75um cameras (Basler AG). The four polarization-resolved paths are constructed using a non-polarizing 50/50 beamsplitter (21014-UF2, Chroma Technology Corp.) and two wire-grid polarizing beamsplitters (FBF04C-UF, Moxtek Inc.). These beamsplitters were chosen for their low wavefront distortions and relatively uniform polarization performance across the visible spectrum. An achromatic half-wave plate (AH-100S-VIS, Meadowlark Optics, Inc.) oriented at −22.5° is placed after the reflection from the 50/50 beamsplitter to create imaging paths sensitive to light polarized along 45° and 135°. A LPVISE100 linear polarizer (Thorabs, Inc.) is placed in front of the I90 and I135 cameras to reject a small p-polarized reflection from the wire-grid beamsplitters.

For label-free imaging we use a 16-channel pE-4000 LED light source (CoolLED Ltd.). The sample is illuminated using a custom-built condenser aligned in Köhler illumination. The condenser uses a changeable 0.4 NA (air) or 1.4 NA (oil immersion) front lens (11525409 or 11551004, Leica Microsystems, Inc.); an aperture stop is used to control the NA of illumination. A broadband circular polarizer constructed from a wire-grid linear polarizer (WP25M-UB, Thorlabs, Inc.) and an achromatic quater-wave plate (AQWP10M-580, Thorlabs, Inc.) is placed in the back focal plane of the condenser lens to illuminate the sample with light of right-hard circular polarization. Epi-fluorescence imaging is carried out with a 7-channel nĳi LED light source (Bluebox Optics Ltd.). We use a non-polarizing 10/90 beamsplitter (21016, Chroma Technology Corp.) to reflect the fluorescence excitation light toward the objective, as dichroic beamsplitter commonly exhibit high diattenuation. Bandpass excitation filters from a Pinkel filter set (LED-DA/FI/TR/Cy5-4X-B-000, Semrock Inc.) are installed in the nĳi light source and a matching quad-band emission filter is used to collect fluorescence emission from multiple channels, unless otherwise specified.

### Microscope automation through hardware sequencing

Microscope control and data acquisition were carried out using Micro-Manager (https://micro-manager.org/) and pycromanager (https://pypi.org/project/pycromanager/). Acquisition from the four cameras was coordinated using the Micro-Manager Multi-Camera adaptor. The exposure signal of the main camera was connected to an Arduino Uno controller and served as a master clock for TTL control of the secondary cameras, the *z* piezo stage (PZ-2000 with 300 um range *z* axis, Applied Scientific Instrumentation), and the light sources (fig. 1c). The secondary cameras were triggered at the master clock rate. The piezo *z* stage was programmed in Micro-Manager for Sequence acquisition and was also triggered at the master clock rate. For control of the light sources, Arduino Shutters were constructed in Micro-Manager and the corresponding shutter pins were connected to the TTL input of the light sources. Multichannel acquisition was carried out through the Micro-Manager Multi-Dimensional Acquisition dialog. For fast single channel acquisition a custom piezo *z* step sequence was constructed in python and uploaded to the stage using pycromanager. The piezo stage was programmed for FastSequence acquisition and a pycromanager event sequence describing the time and *z* dimensions of the acquisition was constructed and sent to the acquisition engine. An example script for fast single channel acquisition is provided here.

### Calibration of polarization response

The instrument tensor is calibrated by illuminating at the label-free imaging wavelength or at the fluorophore emission wavelength and measuring the detected intensity in the four polarization channels of the microscopes for a variety of Stokes states (fig. 2a), realized by rotating a wire-grid linear polarizer (WP25M-UB, Thorlabs, Inc.) in the sample plane with a K10CR1 rotation stage (Thorlabs, Inc.). The Stokes state of the light source, *S*_*illum*_, and the initial angle of the linear polarizer *θ*_0_ are calibrated from a fit to the I0 intensity as a function of polarizer angle *θ*:

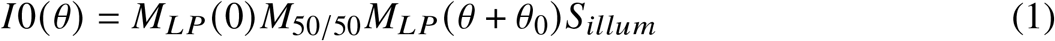

where *M*_*LP*_ *(θ*) and *M*_50/50_ Mueller matrices of a linear polarizer at angle *θ* and a linear diattenuator with 50% transmission. The instrument tensor is then calculated from the Stokes state in the sample plane *S*_*samp*_ (*θ*) = *M*_*LP*_ (*θ* + *θ* _0_)*S*_*illum*_ and the measured intensities in four channels at every pixel *1* (*r*) as:

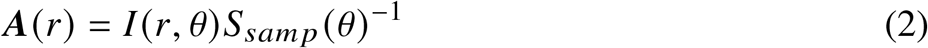

where *A(r)* and *1(r)* are 4*x*3 and 4*x N* matrices, respectively, at pixel position *r, S*_*samp*_ is a 3*x N* matrix, and *N* is the number of angular samples.

### Sample preparation and imaging

Live cell experiments were carried out in an Okolab stage-top environmental chamber set to maintain 37°C and 5% carbon dioxide. All cells types were plated on Cellvis glass bottom well plates for imaging, unless otherwise specified.

U2OS cells were cultured in McCoy’s 5A medium (Thermo Fisher Scientific, 16600108) supplemented with 10 % fetal bovine serum (FBS), 1x penicillin and streptomycin, and 4 mM L-glutamine. For imaging, the media was changed to phenol-red free DMEM (Thermo Fisher Scientific, 31053028) supplemented in the same manner. Cells were stained with 1 µM SiR-actin and 10 µM verapamil (Cytoskeleton, Inc., CY-SC001) for 1-3 hours before imaging. The sample was illuminated with 525 nm light at 0.4 NA for label-free imaging and with 630 nm light for fluorescence imaging. 8 µm z-stacks with 0.25 µm step size were acquired every 30 seconds. Data are presented in fig. 1d-e and accompanying visualizations.

Cardiomyocytes were derived from iPS cells and cultured in RPMI 1640 media supplemented with B27 and Antibiotic-Antimycotic (Thermo Fisher Scientific, 11875093, 17504044, and 15240062), as previously described [17]. For experiments shown in fig. 4b cells were treated with 1 µM SiR-actin and 10 µM verapamil for 2 hours. The SiR-actin staining media was aspirated, and Hoechst-33342 (Thermo Fisher Scientific, 62249) was added at 1 ug/mL in culture media for 30 minutes. Cells were washed 1x with PBS and culture media was replaced before imaging. For experiments shown in fig. 3 the cell culture media was replaced 1-2 hours before imaging with RPMI media (Sigma-Aldric R8755) supplemented with B27, Antibiotic-Antimycotic, 2 g/L sodium bicarbonate, 25 mM HEPES, and 20% OptiPrep (Sigma-Aldric, D1556) at pH 7.3. We varied the OptiPrep concentration between 0% and 50% when optimizing sarcomere imaging conditions (fig. S7) [39]. Cells were imaged under 525 nm illumination at 0.4 NA. Data in fig. 3a and fig. S7 was acquired in a single plane at 20 Hz. Data in fig. 3e was acquired using the custom triangle *z*-axis waveform shown in fig. 1c at reduced field of view of 500 × 600 pixels. 5 µm z-stacks with 0.25 µm step size were acquired at 10 Hz for 10 seconds.

HEK293T cells expressing CAAX-mScarlett were cultured and imaged in supplemented DMEM media. For imaging cells were grown in custom made strain-free chambers made from two glass coverslips with 0.8 mm spacers. Cells were stained with SiR-actin as described above. The sample was illuminated with 490 nm light at 0.8 NA using an oil immersion condenser for label-free imaging and with 545 nm and 630 nm light for fluorescence imaging. 20 µm z-stacks with 0.25 µm step size were acquired every 45 seconds. Data are presented in fig. 4a and accompanying visualizations.

Mouse brain tissue sections were processed as previously described [17]. The corpus callosum region of a 12 µm thick slice was imaged using a Nikon 10x 0.45 NA Plan Apo objective and 0.4 NA illumination. 20 µm z-stacks with 1 µm step size were acquired at eight different wavelengths (see fig. 4c).

### Reconstruction of density, anisotropy, and orientation

Raw intensity images collected by the four cameras were registered using the SURF image-based registration algorithm in MATLAB [59]. Registered images were multiplied by the inverse instrument tensor to calculate Stokes images. Transmission, 3D phase, retardance, and orientation were computed from Stokes images as previously described [16, 17]. Fluorescence anisotropy and orientation were also computed as previously described [10]. Fluorescence intensity was deconvolved with Tikhonov regularization and positivity constraint as summarized in [60]. The data reconstruction pipelines are illustrated with examples in figs. S5 and S6.

Perceptually uniform color overlays of retardance and orientation or fluorescence intensity, anisotropy, and orientation were built using the colorspacious package https://colorspacious.readthedocs.io/en/latest/. All steps of the reconstruction pipeline apart from image registration were carried out in python.

### Denoising and background correction of label-free data

For label-free data reconstruction, Stokes images were denoised using a soft-threshold wavelet denoising algorithm [61]. Anisotropy measurements from U2OS cells (fig. 1e) were processed using sample background correction followed by surface fit background correction. Stokes parameters were averaged along *z* in chunks of four before computing retardance and orientation. Data of cardiomyocyte beating dynamics (fig. 3a) were corrected using background images constructed by averaging nine fields of view. Averaging multiple sample Stokes images removes the anisotropic contribution of the sample but preserved background signal present in all images; this background correction technique is useful when an empty area within the sample is challenging to find.

For mouse brain data (fig. 4c) we reconstructed 2D phase and projected retardance values along *z* before taking average measurements of physical properties at different wavelengths over a region of interest.

### Denoising and background correction of fluorescence data

For fluorescence anisotropy reconstruction, intensity images were denoised before conversion to Stokes space using the same algorithm as above. We assumed that the fluorescence excitation source is unpolarized and used background correction methods to account for polarization distortions in fluorescence anisotropy measurements. Anisotropy measurements of actin in U2OS cells(fig. 1d) and of DNA and actin in cardiomyocytes (fig. 4b) were background-corrected using the median of *S*1_*norm*_ and *S*2_*norm*_ channels calculated over six and three fields of view, respectively. Assuming that background fluorescence polarization effects are slowly varying in space, we carried out background correction on fluorescence anisotropy measurements of CAAX-stained membrane and SiR-actin-stained F-actin in HEK293T cells (fig. 4a) using uniform-filtered *S*1_*raw*_ and *S*2_*raw*_ channels with filter kernel size much larger than the size of cells (in this case, 200 ×200 ×20 in XYZ pixel space). Calibration methods accounting for partial polarization of the fluorescence excitation source are described in [10].

### Optical flow analysis

Sparse and dense optical flow analysis of motion in cardiomyocytes was carried out using the Lucas-Kanade and Farneback algorithms, respectively, as implemented in OpenCV [40, 41, 62]. Displacement and velocity traces were reconstructed from *XY* coordinates of points of interest over time.

## Supporting information

Visualization 1

Visualization 2

Visualization 3

Visualization 4

Visualization 5

Visualization 6

Visualization 7

Visualization 8

Visualization 9

Visualization 10

Visualization 11

Visualization 12

Visualization 13

Visualization 14

Visualization 15

Visualization 16

## Acknowledgments

We thank the May Han lab at Stanford University for providing mouse brain tissue slices. We thank Cameron Foltz for contributing to the microscope automation and data management tools. We also thank our past lab members Jaehee Park for contributing to the phase reconstruction and calibration algorithms, and Elliot Mount for contributing to the initial build of the microscope.

## Data availability

Raw images and microscope calibration metadata for reconstruction of data presented in fig. 1d-e and fig. 3e are available at https://doi.org/10.5281/zenodo.5952953. Jupyter notebooks outlining steps in the reconstruction process are available at https://github.com/mehta-lab/miPolScope.

## Supplemental document

See Supplement 1 for supporting content.

