## Supplement 1 for "Correlative imaging of spatio-angular dynamics of molecular assemblies and cells with multimodal instant polarization microscope"

### 1. SUPPLEMENTAL FIGURES

**(a)**

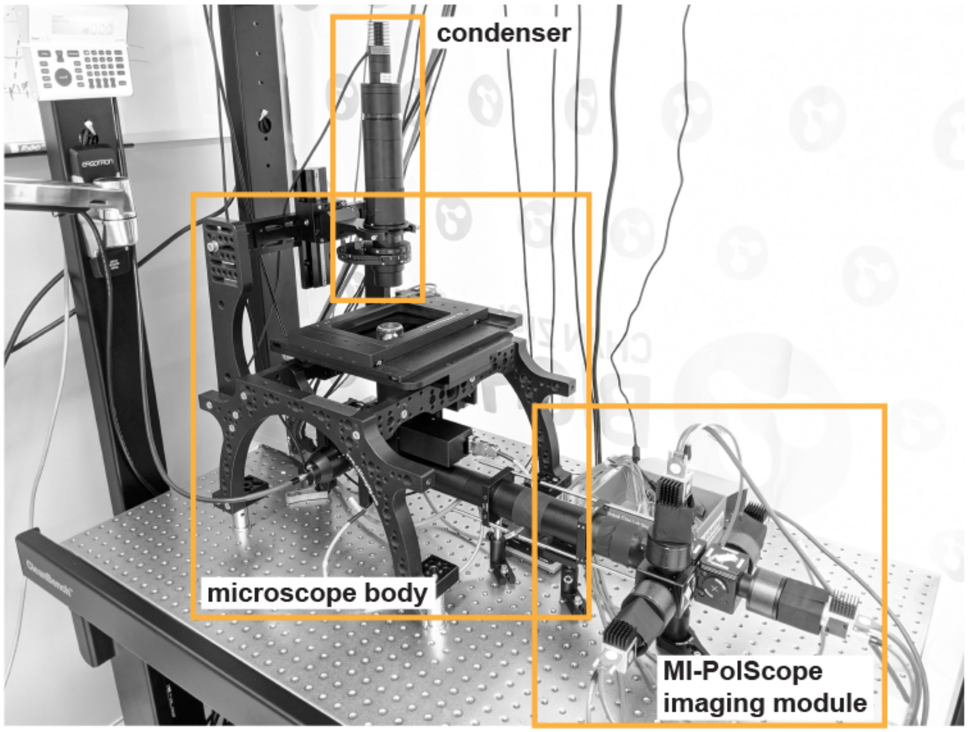


**(b)**

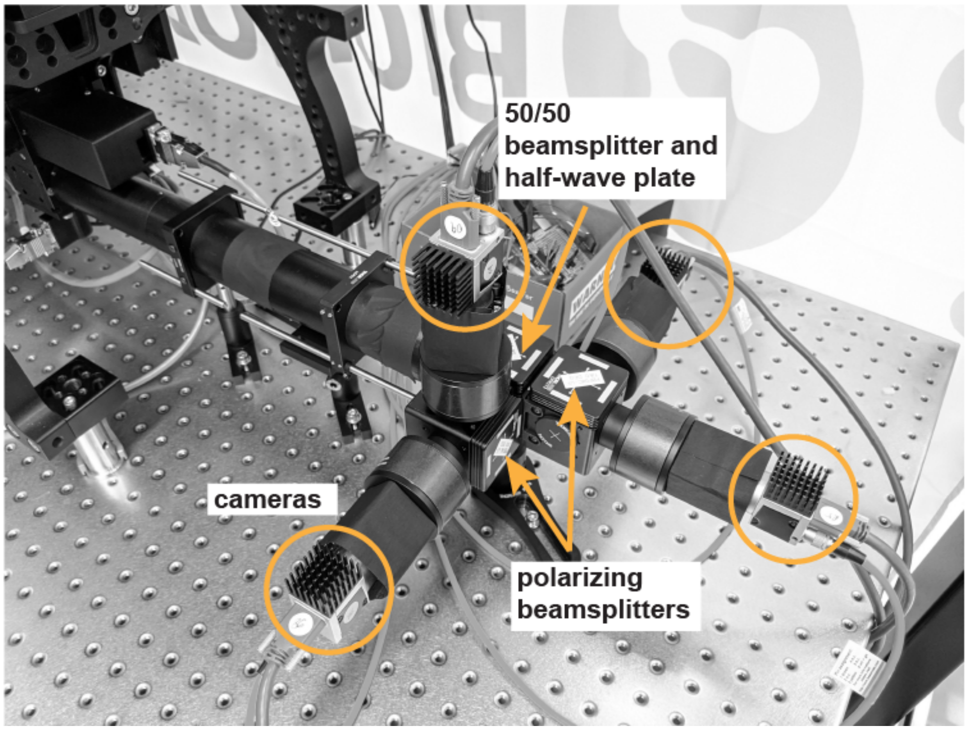


**Fig. S1. Pictures of the microscope (a) and the miPolScope imaging module (b).** Key components of the microscope are highlighted in orange.

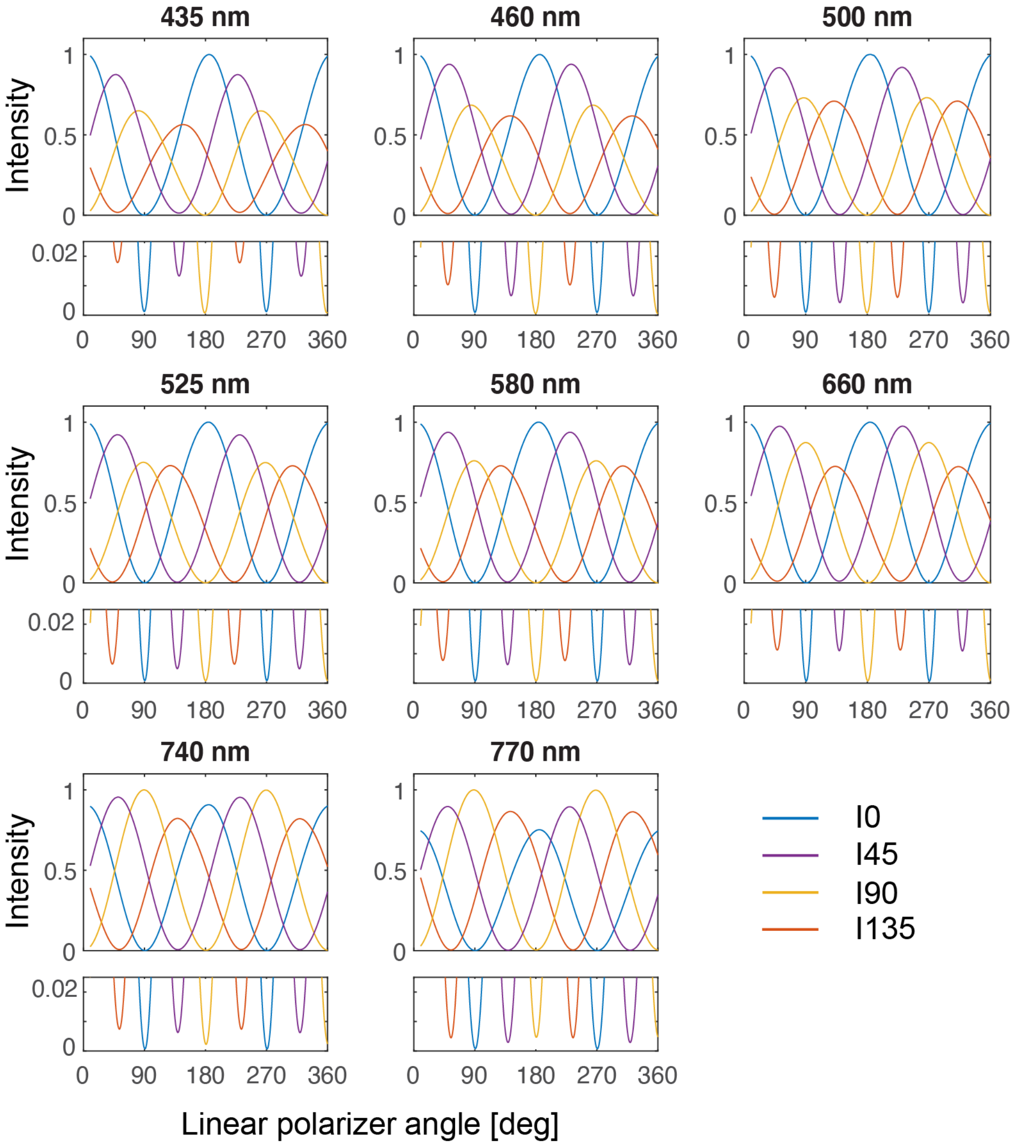


**Fig. S2. Measured intensity in four channels over a central region of the microscope field of view at eight different wavelengths.** The extinction ratio in each channel is given in table S1. The extinction ratio in I45 and I135 is reduced due to small deviation from the theoretical retardance of the achromatic half-wave plate.

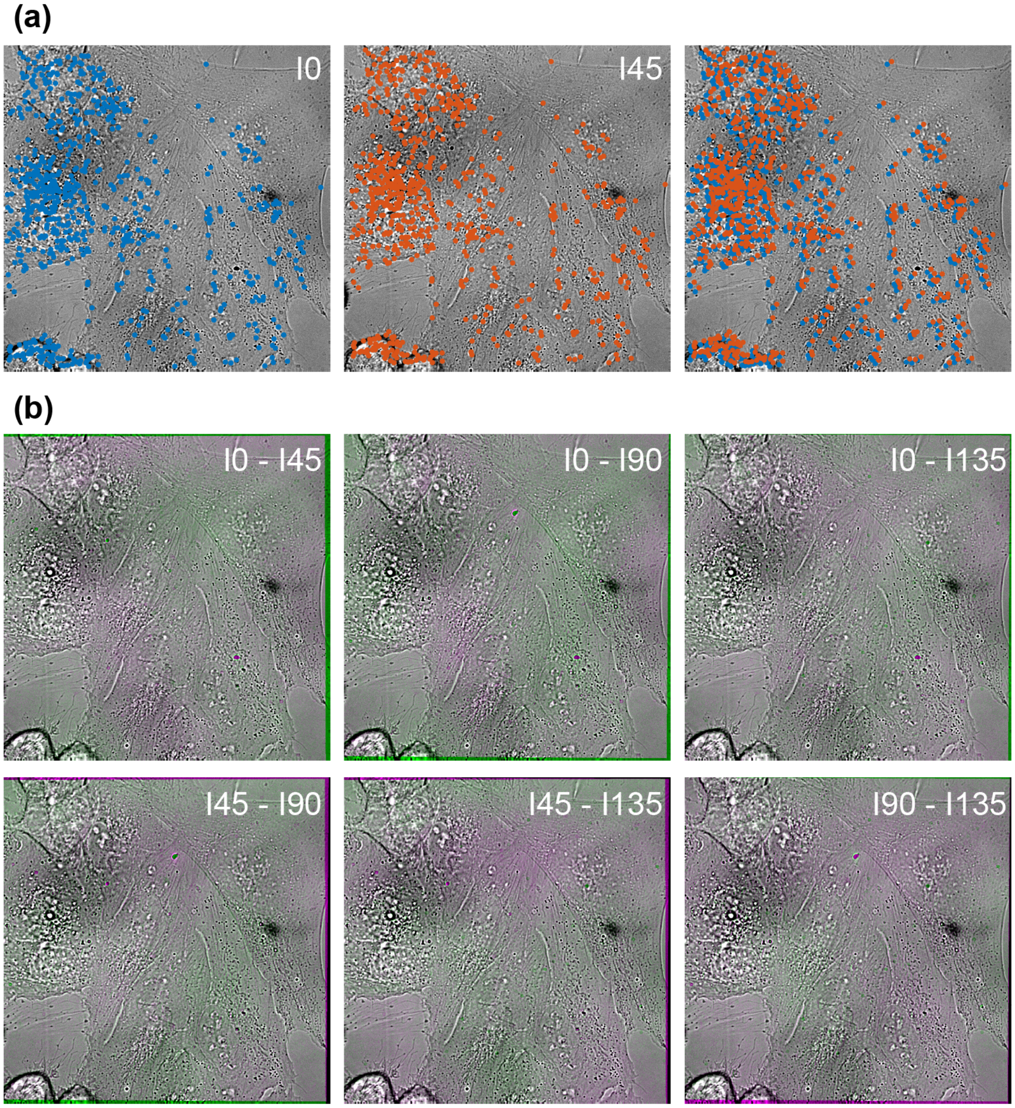


**Fig. S3. Registration of images from the four detectors of the miPolScope imaging module**. Raw images were registered using a projective geometric transformation calculated based on the SURF image-based registration algorithm in MATLAB [1]. **(a)** Left and middle: Detected features in the I0 and I45 channels are shown as blue and orange dots. Overlay of the features (right) highlights misregistration between the two channels. **(b)** Green-magenta pairwise overlays of the four channels after image registration. Images have been flat-field corrected with a fitted 2D surface for display.

(a)

(b)

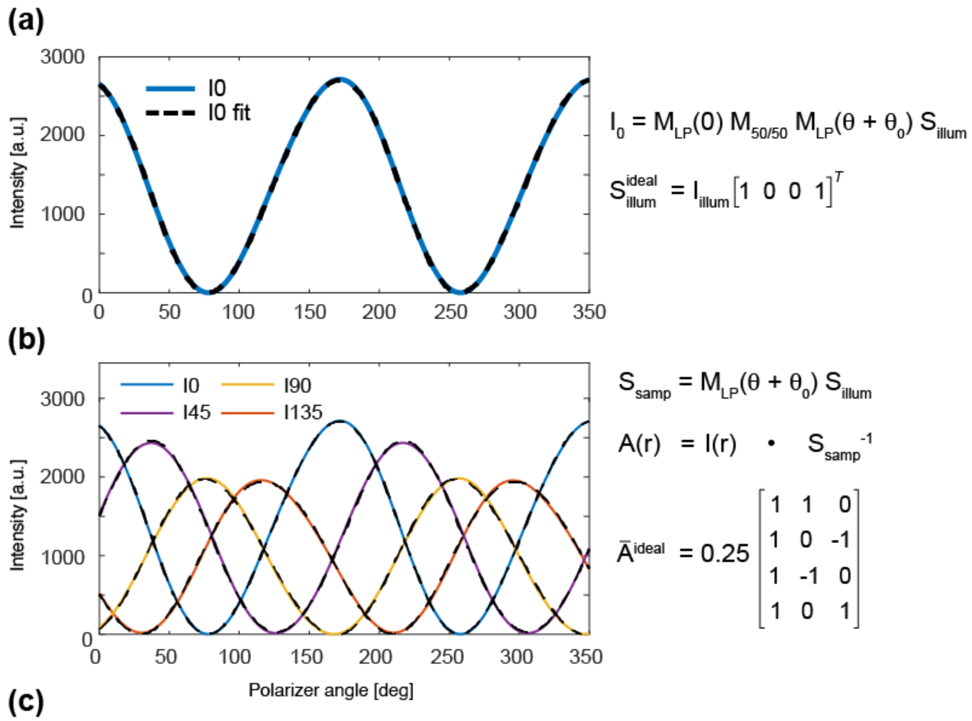


(c)

| Wavelength | $435\ nm$ | $460\ nm$ | $500\ nm$ | $525\ nm$ |
|---|---|---|---|---|
| $S_{illum,\lambda}$ | $\begin{bmatrix} 1 \\ 0.029 \\ 0.126 \end{bmatrix}$ | $\begin{bmatrix} 1 \\ -0.004 \\ 0.102 \end{bmatrix}$ | $\begin{bmatrix} 1 \\ -0.030 \\ 0.074 \end{bmatrix}$ | $\begin{bmatrix} 1 \\ -0.050 \\ 0.064 \end{bmatrix}$ |
| $\bar{A}_\lambda$ | $\begin{bmatrix} 0.254 & 0.252 & -0.002 \\ 0.175 & 0.028 & -0.165 \\ 0.186 & -0.189 & -0.002 \\ 0.218 & -0.051 & 0.200 \end{bmatrix}$ | $\begin{bmatrix} 0.279 & 0.277 & 0.000 \\ 0.202 & 0.031 & -0.195 \\ 0.197 & -0.199 & -0.006 \\ 0.259 & -0.079 & 0.236 \end{bmatrix}$ | $\begin{bmatrix} 0.275 & 0.274 & -0.010 \\ 0.208 & -0.004 & -0.208 \\ 0.191 & -0.191 & -0.011 \\ 0.232 & -0.043 & 0.220 \end{bmatrix}$ | $\begin{bmatrix} 0.279 & 0.279 & -0.001 \\ 0.207 & -0.014 & -0.208 \\ 0.190 & -0.188 & -0.013 \\ 0.234 & -0.027 & 0.224 \end{bmatrix}$ |

| Wavelength | $580\ nm$ | $660\ nm$ | $740\ nm$ | $770\ nm$ |
|---|---|---|---|---|
| $S_{illum,\lambda}$ | $\begin{bmatrix} 1 \\ -0.055 \\ 0.059 \end{bmatrix}$ | $\begin{bmatrix} 1 \\ -0.075 \\ 0.051 \end{bmatrix}$ | $\begin{bmatrix} 1 \\ -0.074 \\ 0.059 \end{bmatrix}$ | $\begin{bmatrix} 1 \\ -0.070 \\ 0.072 \end{bmatrix}$ |
| $\bar{A}_\lambda$ | $\begin{bmatrix} 0.289 & 0.289 & 0.000 \\ 0.211 & -0.013 & -0.211 \\ 0.196 & -0.194 & -0.015 \\ 0.253 & -0.026 & 0.242 \end{bmatrix}$ | $\begin{bmatrix} 0.264 & 0.265 & -0.001 \\ 0.190 & 0.026 & -0.188 \\ 0.200 & -0.195 & -0.018 \\ 0.237 & -0.027 & 0.223 \end{bmatrix}$ | $\begin{bmatrix} 0.243 & 0.244 & -0.001 \\ 0.228 & -0.066 & -0.220 \\ 0.242 & -0.234 & -0.020 \\ 0.241 & -0.028 & 0.231 \end{bmatrix}$ | $\begin{bmatrix} 0.214 & 0.215 & -0.001 \\ 0.260 & 0.091 & -0.247 \\ 0.262 & -0.252 & -0.020 \\ 0.256 & -0.025 & 0.229 \end{bmatrix}$ |

**Fig. S4. Calibration of the Mi-PolScope illumination and detection.** Microscope calibration is carried out by rotating a linear polarizer in the sample plane as described in *Methods* and fig. 2a. **(a)** The Stokes state of the illumination light $S_{illum}$ is calculated from a fit of the intensity in the I0 channels to the model $I0(\theta) = M_{LP}(0)M_{50/50}M_{LP}(\theta + \theta_0)S_{illum}$. The ideal Stokes state of illumination is that of right-hand circularly polarized light with intensity $I_{illum}$. **(b)** The instrument tensor for the microscope, $\boldsymbol{A}$, is calculated as the least-squares solution of $\boldsymbol{A}(r) = I(r,\theta)S_{samp}(\theta)^{-1}$ where $I(r)$ is the intensity in four channels for every pixel $r$ and $S_{samp}(\theta) = M_{LP}(\theta + \theta_0)S_{illum}$ is the Stokes state of light in the sample plane, i.e. after the rotating linear polarizer. The plot on the left shows measured intensity in four channels (solid colored lines) and intensities calculated based on the calibrated instrument tensor $I^{cal} = AS_{samp}$ (dashed black lines). The ideal instrument matrix for the microscope ($\bar{A}$, i.e. instrument tensor averaged over the image dimensions) is shown on the right. **(c)** Measurements of the Stokes state of illumination and instrument matrix at eight different wavelengths.

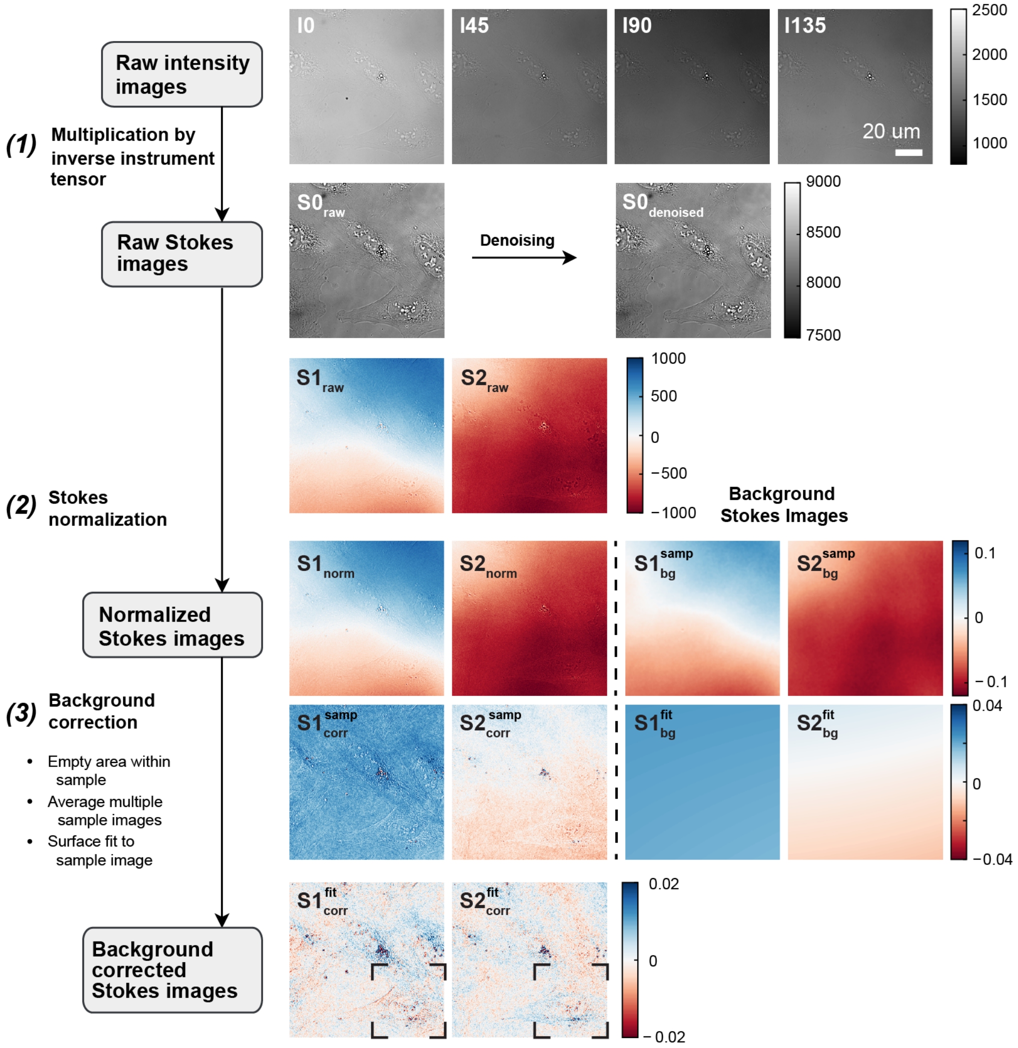


**Fig. S5. Flowchart showing steps in label-free data reconstruction.** Raw intensity images are multiplied by the inverse instrument tensor **(1)** to obtain raw Stokes images. $S0_{raw}$ is denoised using wavelet denoising. $S1_{raw}$ and $S2_{raw}$ are normalized **(2)** and background corrected **(3)**. Background correction removes the contribution of birefringence of the sample chamber and may be carried out using: (i) Stokes images collected at an empty area of the sample, (ii) Stokes images constructed by averaging multiple sample images, (iii) 2D surface fit to the sample Stokes images, or a combination of these. In this example, normalized Stokes images are corrected by subtracting a sample background ($S1_{bg}^{samp}$ and $S2_{bg}^{samp}$, shown on the right) followed by 2D surface fit to the residuals ($S1_{bg}^{fit}$ and $S2_{bg}^{fit}$) to remove slow-varying components. Corrected Stokes images are then denoised. Black rectangles show regions of interest used in the next panel.

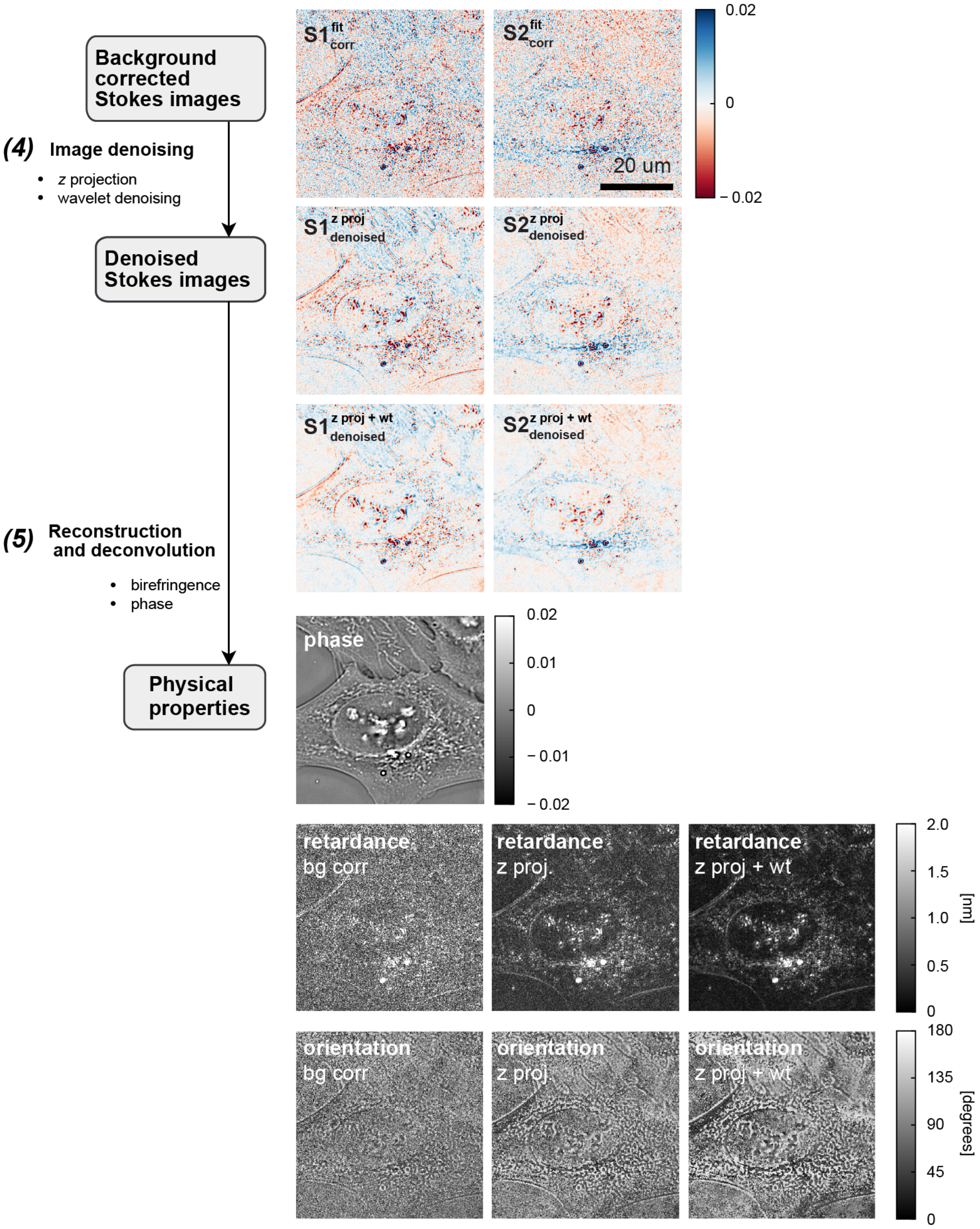


**Fig. S5.** (cont.) Corrected Stokes images are denoised **(4)** using: (i) mean projection of multiple z slices, (ii) wavelet denoising, or a combination of the two. In this example, we show zoomed-in regions highlighted in the previous panel. Stokes images are denoised by averaging of blocks of four z-slices ($S1^{z\ proj}_{denoised}$ and $S2^{z\ proj}_{denoised}$) followed by wavelet denoising ($S1^{z\ proj+wt}_{denoised}$ and $S2^{z\ proj+wt}_{denoised}$). Retardance and orientation are reconstructed **(5)** from denoised $S1$ and $S2$ images. For comparison, we show reconstructions using partial data processing - using background correction alone (left, based on $S1^{fit}_{corr}$ and $S2^{fit}_{corr}$), background correction and z-projection (middle, based on $S1^{z\ proj}_{denoised}$ and $S2^{z\ proj}_{denoised}$), and using the full pipeline, on the right. Phase can be reconstructed from raw $S0$ images **(5)** as the reconstruction algorithm has built-in signal regularization.

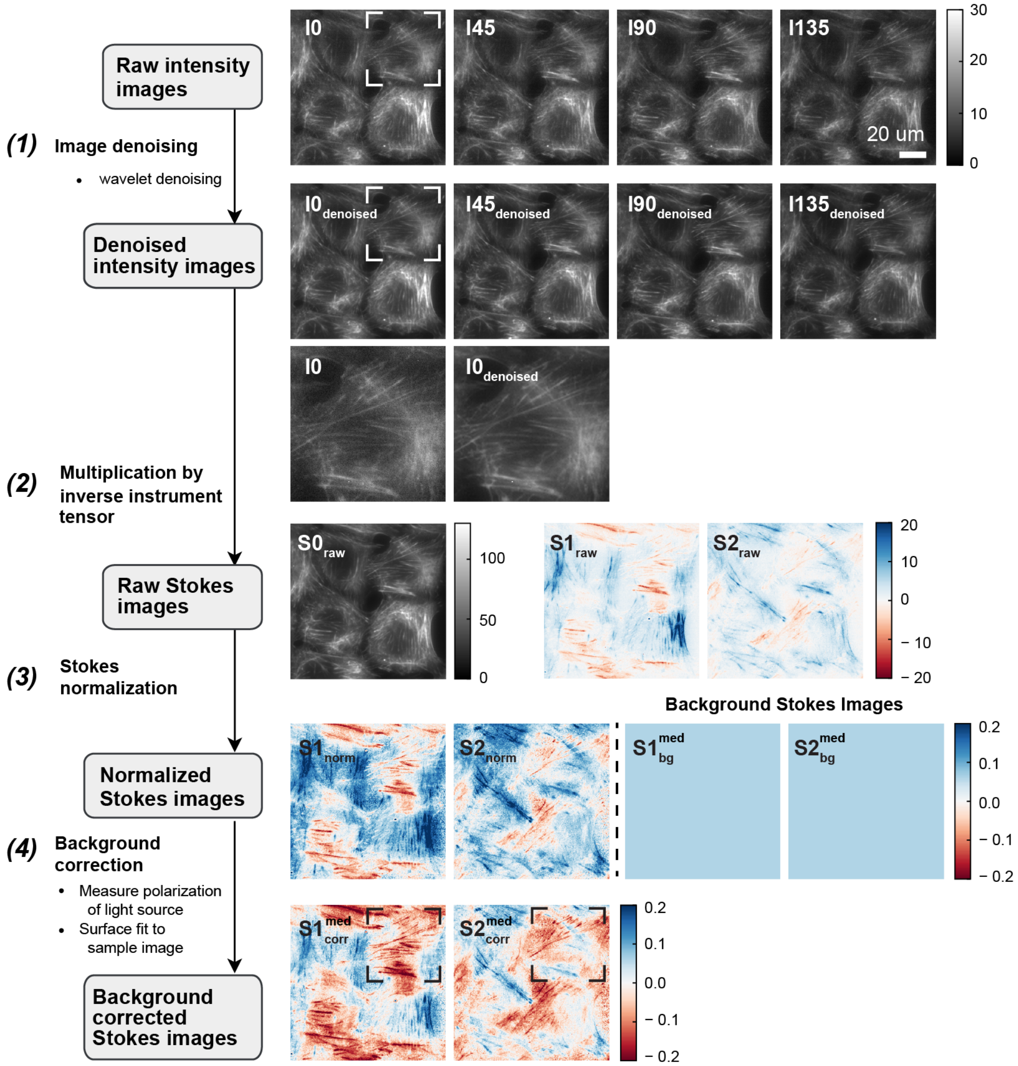


**Fig. S6. Flowchart showing steps in fluorescence anisotropy data reconstruction.** Raw intensity images are denoised **(1)** using wavelet denoising. Highlighted regions in $I0$ and $I0_{denoised}$ are enlarged to show the effect of image denoising. Denoised images are multiplied by the inverse instrument tensor **(2)** to obtain raw Stokes images. $S1_{raw}$ and $S2_{raw}$ are normalized **(3)** and background corrected **(4)**. Background correction removes bias due to partial polarization of the fluorescence excitation source and may be carried out by: (i) measuring the polarization state of the excitation source (e.g. using a sample of static randomly oriented fluorophores such as a dyed plastic slide), (ii) fitting a 2D surface to the sample Stokes images (provided that structures are isotropically oriented at large scales), or using a combination of the two. In this example, normalized Stokes images are corrected by subtracting the median value of each image ($S1_{bg}^{med}$ and $S2_{bg}^{med}$, shown as constant-value surfaces on the right). Corrected $S1$ and $S2$ images are then used for reconstruction of fluorescence anisotropy and orientation. Black rectangles show regions of interest used in the next panel.

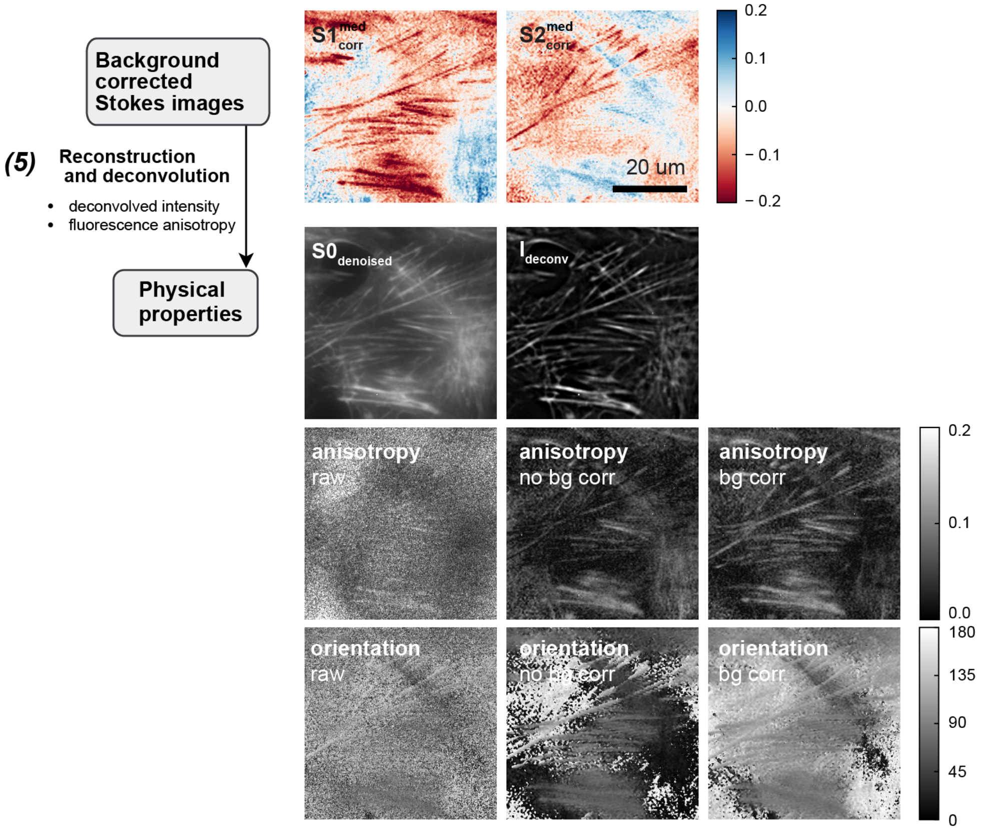


**Fig. S6.** (cont.) Zoomed-in regions of corrected $S1$ and $S2$ images highlighted in the previous panel are shown. Raw $S0$ images are used for deconvolution of fluorescence intensity **(5)**. For comparison, we show denoised S0 and deconvolved fluorescence intensity images. Corrected $S1$ and $S2$ images are used for reconstruction of fluorescence anisotropy and orientation **(5)**. For comparison, we show reconstructions using partial data processing - without image denoising (left, based on $I0$, $I45$, $I90$, and $I135$), without background correction (middle, based on $S1_{norm}$ and $S2_{norm}$), and using the full pipeline, on the right.

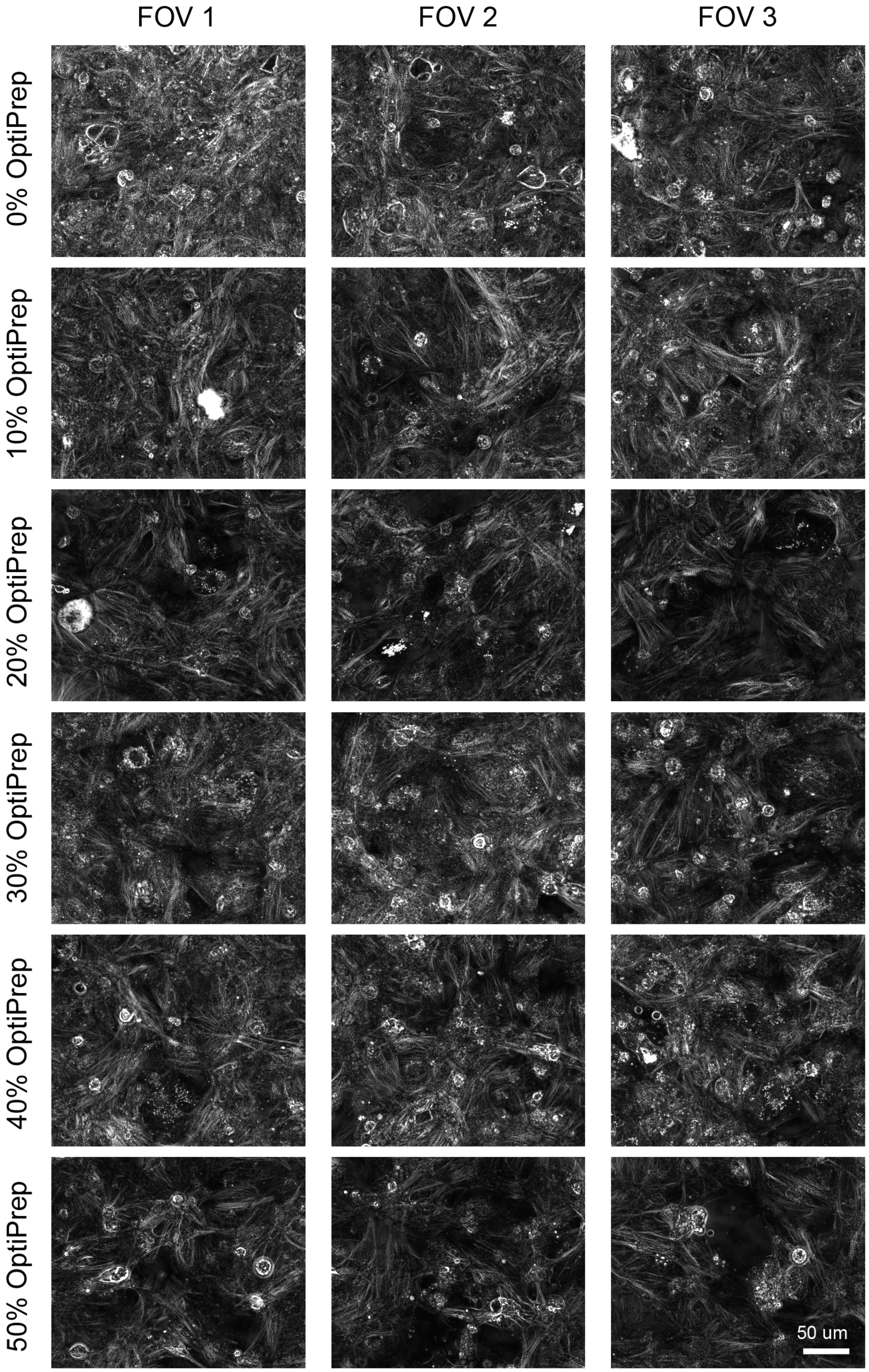


**Fig. S7. Retardance measurements in live cardiomyocytes using media with varying fraction of the OptiPrep index matching reagent.** Based on the fraction of OptiPrep in the media we estimated that the refractive index varies between 1.34 (0% OptiPrep) and 1.42 (50% OptiPrep). We found that media with 20% OptiPrep (refractive index of 1.37) enhances the retardance contrast from sarcomeres.

### 2. SUPPLEMENTAL TABLES

| | 435 nm | 460 nm | 500 nm | 525 nm | 580 nm | 660 nm | 740 nm | 770 nm |
|---|---|---|---|---|---|---|---|---|
| I0 | 950:1 | 1200:1 | 1460:1 | 2090:1 | 2170:1 | 2740:1 | 2940:1 | 2210:1 |
| I45 | 30:1 | 50:1 | 110:1 | 100:1 | 90:1 | 60:1 | 100:1 | 190:1 |
| I90 | 1310:1 | 2230:1 | 1630:1 | 1950:1 | 1830:1 | 2220:1 | 550:1 | 240:1 |
| I135 | 60:1 | 120:1 | 200:1 | 170:1 | 130:1 | 80:1 | 150:1 | 330:1 |

**Table S1.** Extinction ratio in four channels as a function of wavelength.

| $I0 \leftarrow I45$ | $I0 \leftarrow I90$ | $I0 \leftarrow I135$ |
|---|---|---|
| $\begin{bmatrix} 1.0014 & -0.0034 & 0 \\ 0.0039 & 1.0022 & 0 \\ -14.53 & 10.75 & 1 \end{bmatrix}$ | $\begin{bmatrix} 1.0018 & 0.0071 & 0 \\ -0.0065 & 1.0034 & 0 \\ -1.74 & -12.02 & 1 \end{bmatrix}$ | $\begin{bmatrix} 1.0007 & 0.0045 & 0 \\ -0.0046 & 1.0006 & 0 \\ -0.92 & 2.28 & 1 \end{bmatrix}$ |

**Table S2.** Transformation matrices, $T$, for registration of images from four cameras. Images from $I45$, $I90$, and $I135$ channels are registered to the reference $I0$ channel. Transformation matrices are computed for every dataset, and as necessary, for every field of view. Points in the old coordinate system $(x, y)$ are transformed into the new coordinate system $(u, v)$ as $[u\ v] = [x\ y\ 1]T$.

### 3. LIST OF VIDEOS

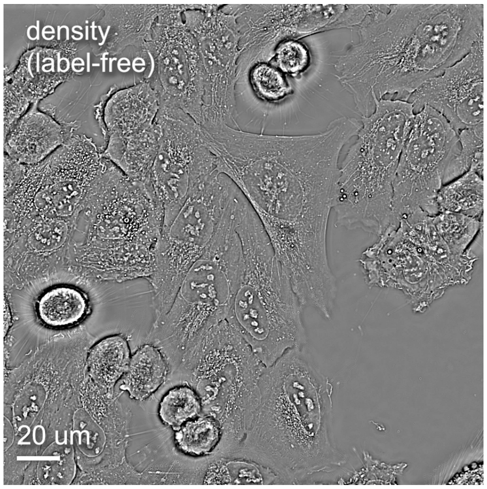


**Visualization 1.** Overview of density and anisotropy measurements of U2OS cells imaged over time in label-free and fluorescence modes. Overlays of phase & retardance and phase & fluorescence anisotropy highlight complementary information obtained from different channels.

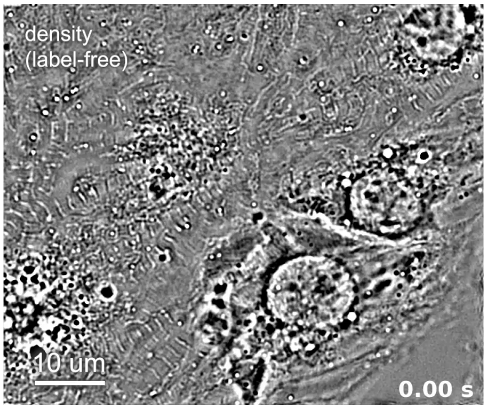


**Visualization 2.** Overview of density and anisotropy measurements of live cardiomyocytes. Complementary modes of information highlight distinct structures in myofibrils. High-speed imaging enables detailed analysis of contractile dynamics.

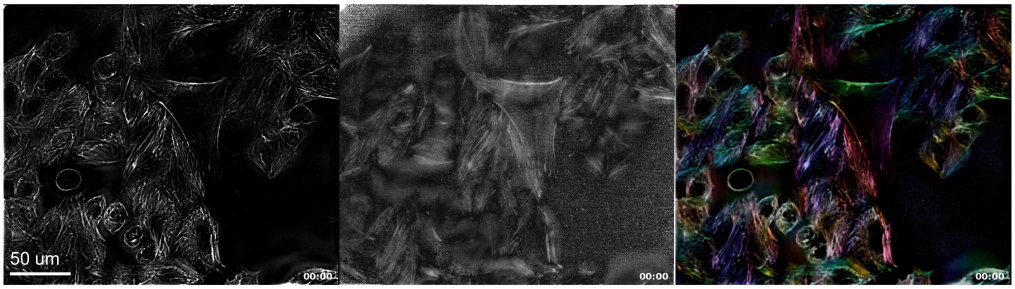


**Visualization 3.** Time series movie of SiR-actin deconvolved intensity (left), anisotropy (middle), and intensity, anisotropy, and orientation overlay (right) in labeled U2OS cells at full field of view of the microscope.

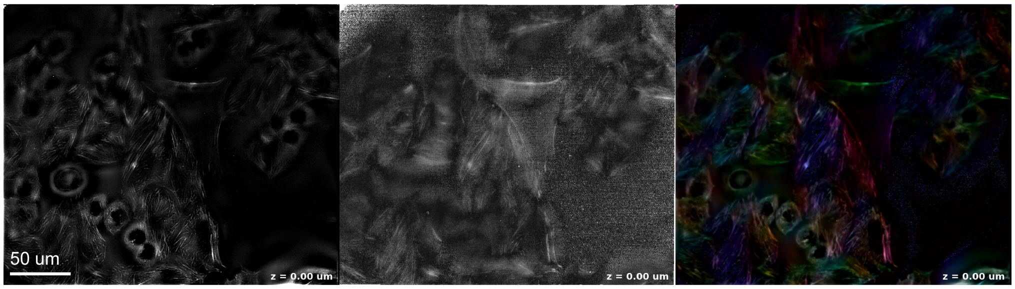


**Visualization 4.** Axial fly-through of SiR-actin deconvolved intensity (left), anisotropy (middle), and intensity, anisotropy, and orientation overlay (right) in labeled U2OS cells at full field of view of the microscope.

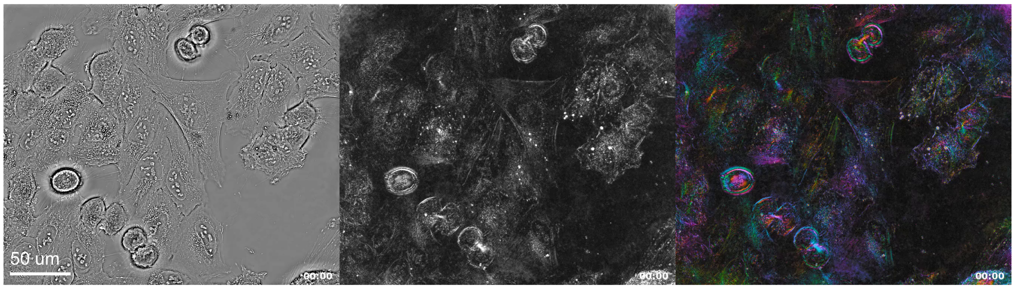


**Visualization 5.** Time series movie of phase (left), retardance (middle), and retardance and orientation overlay (right) in U2OS cells at full field of view of the microscope.

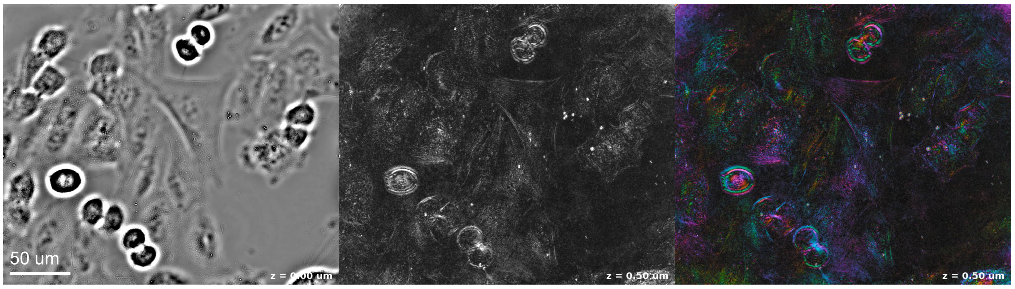


**Visualization 6.** Axial fly-through of phase (left), retardance (middle), and retardance and orientation overlay (right) in U2OS cells at full field of view of the microscope.

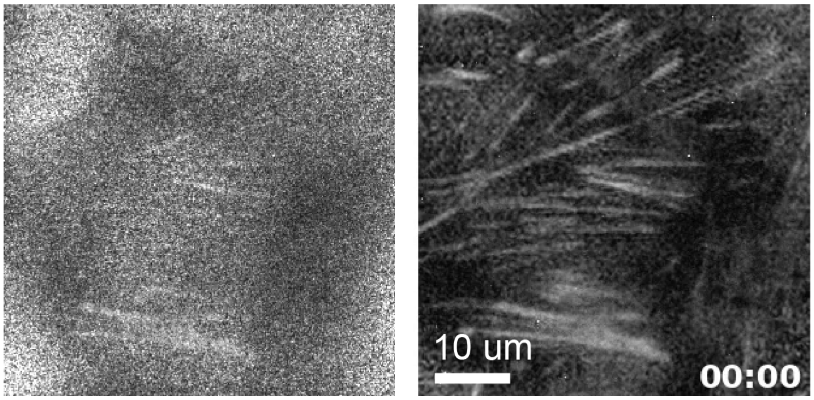


**Visualization 7.** Time series movie of raw (left) and denoised (right) fluorescence anisotropy measurements of labeled actin in U2OS cells.

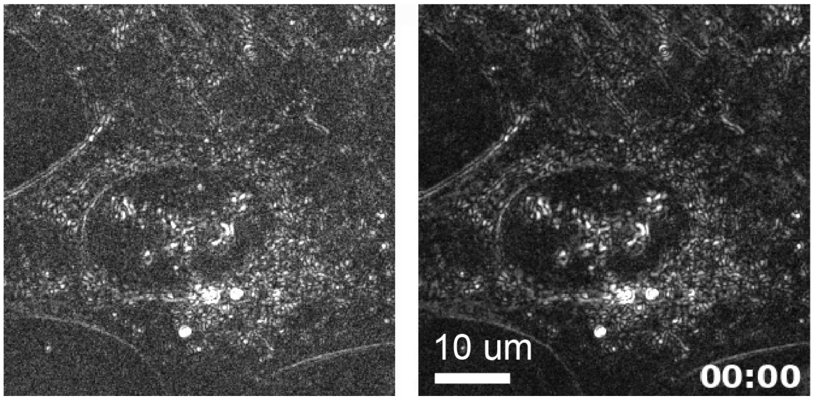


**Visualization 8.** Time series movie of raw (left) and denoised (right) retardance measurements in U2OS cells.

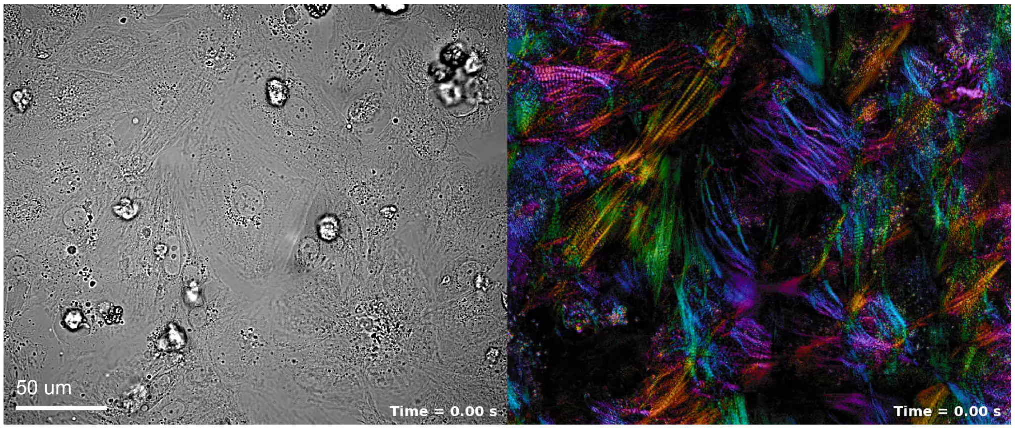


**Visualization 9.** Time series movie of computed brightfield (left) and retardance and orientation overlay (right) in beating iPSC-derived cardiomyocytes at full field of view of the microscope.

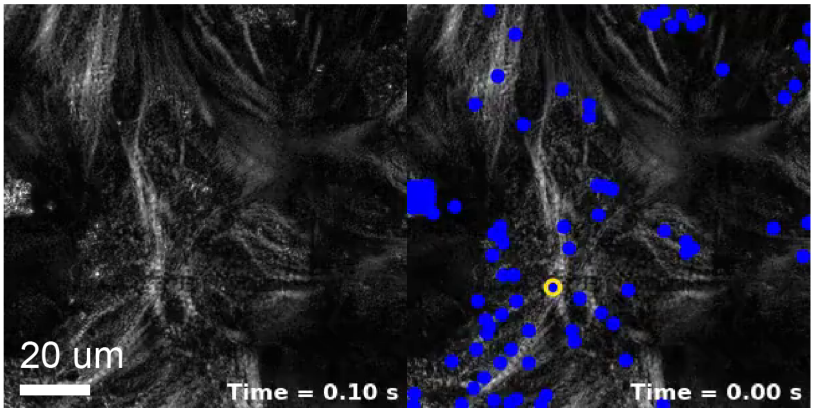


**Visualization 10.** Left: Time series movie of dense optical flow measurements in beating iPSC-derived cardiomyocytes at a region of interest. Displacement is calculated over every two frames of the acquisition. Arrows are plotted for features with displacement greater than 2 pixels. Arrows are 10x longer than the measured displacement. Right: Time series movie of sparse optical flow measurements for the same region of interest.

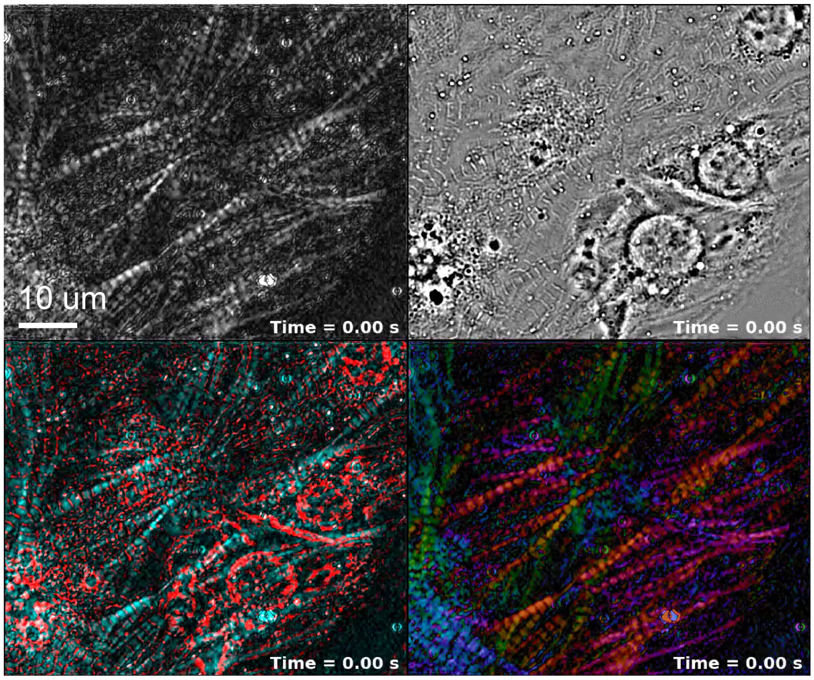


**Visualization 11.** Time series movie of retardance (top left), phase (top right), phase and retardance overlay (bottom left, phase shown in red and retardance shown in cyan), and retardance and orientation overlay (bottom right) in a single plane of a 3D acquisition of beating cardiomyocytes.

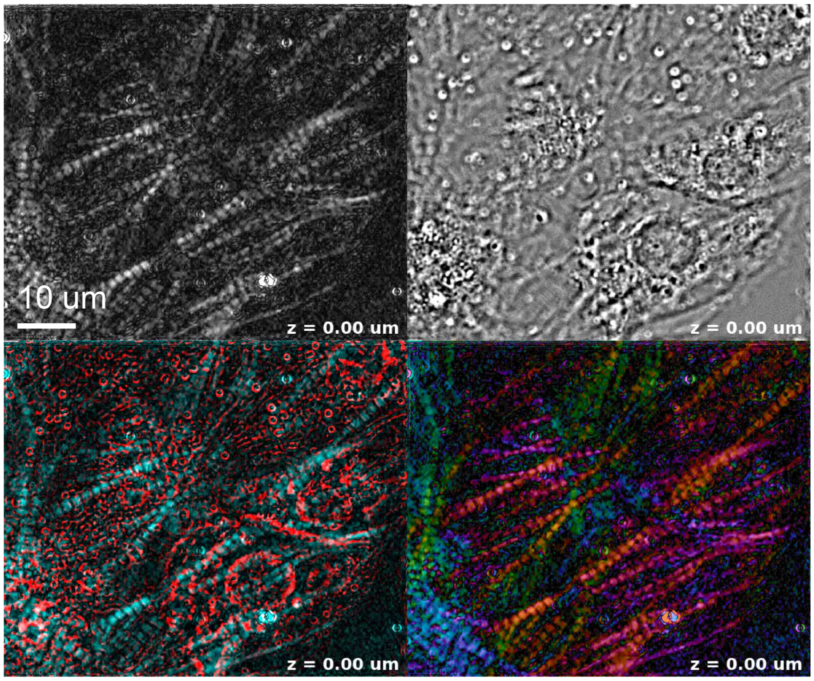


**Visualization 12.** Axial fly-through of retardance (top left), phase (top right), phase and retardance overlay (bottom left, phase shown in red and retardance shown in cyan), and retardance and orientation overlay (bottom right) in beating cardiomyocytes.

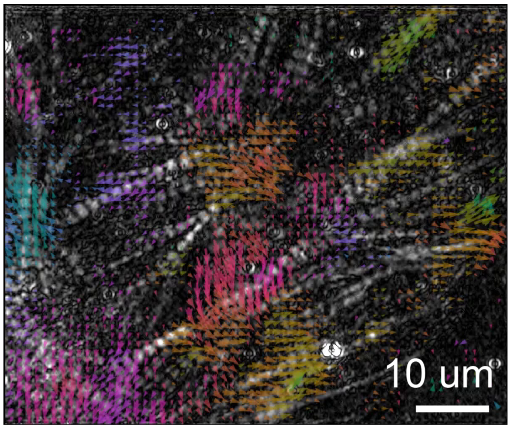


**Visualization 13.** Time series movie of dense optical flow measurements in a single plane in beating cardiomyocytes.

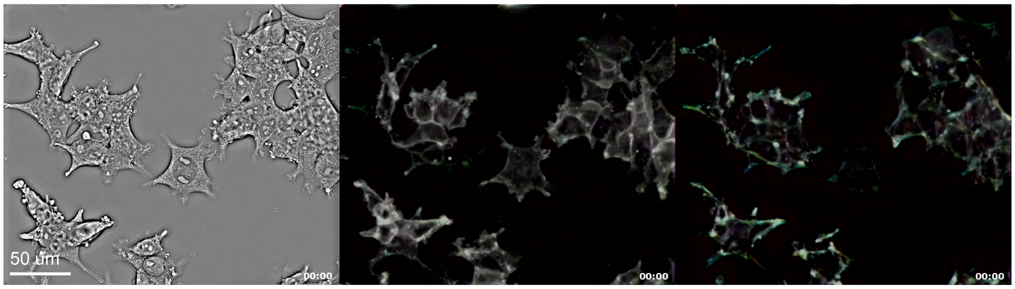


**Visualization 14.** Time series movie of phase (left), CAAX-mScarlett intensity, anisotropy, and orientation overlay (middle), and SiR-actin intensity, anisotropy, and orientation overlay (right) in HEK293-T cells at full field of view of the microscope.

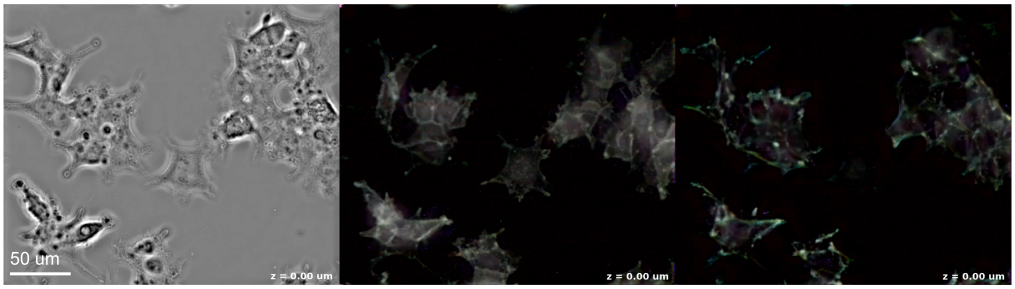


**Visualization 15.** Axial fly-through of phase (left), CAAX-mScarlett intensity, anisotropy, and orientation overlay (middle), and SiR-actin intensity, anisotropy, and orientation overlay (right) in HEK293-T cells at full field of view of the microscope.

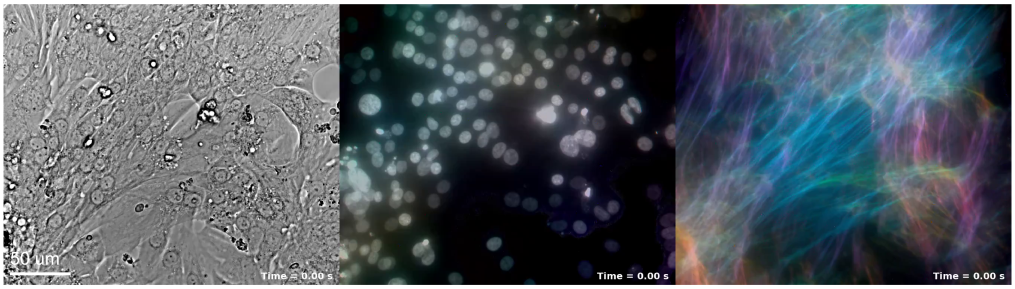


**Visualization 16.** Time series movie of transmission (left), fluorescence anisotropy of Hoechst-labeled DNA (middle), and SiR-actin labeled DNA (right) in beating iPSC-derived cardiomyocytes at full field of view of the microscope.
